# Post-entry, spike-dependent replication advantage of B.1.1.7 and B.1.617.2 over B.1 SARS-CoV-2 in an ACE2-deficient human lung cell line

**DOI:** 10.1101/2021.10.20.465121

**Authors:** Daniela Niemeyer, Simon Schroeder, Kirstin Friedmann, Friderike Weege, Jakob Trimpert, Anja Richter, Saskia Stenzel, Jenny Jansen, Jackson Emanuel, Julia Kazmierski, Fabian Pott, Lara M. Jeworowski, Ruth Olmer, Mark-Christian Jaboreck, Beate Tenner, Jan Papies, Julian Heinze, Felix Walper, Marie L. Schmidt, Nicolas Heinemann, Elisabeth Möncke-Buchner, Talitha Veith, Morris Baumgardt, Karen Hoffmann, Marek Widera, Tran Thi Nhu Thao, Anita Balázs, Jessica Schulze, Christin Mache, Markus Morkel, Sandra Ciesek, Leif G. Hanitsch, Marcus A. Mall, Andreas C. Hocke, Volker Thiel, Klaus Osterrieder, Thorsten Wolff, Ulrich Martin, Victor M. Corman, Marcel A. Müller, Christine Goffinet, Christian Drosten

## Abstract

Epidemiological data demonstrate that SARS-CoV-2 variants of concern (VOC) B.1.1.7 and B.1.617.2 are more transmissible and infections are associated with a higher mortality than non-VOC virus infections. Phenotypic properties underlying their enhanced spread in the human population remain unknown. B.1.1.7 virus isolates displayed inferior or equivalent spread in most cell lines and primary cells compared to an ancestral B.1 SARS-CoV-2, and were outcompeted by the latter. Lower infectivity and delayed entry kinetics of B.1.1.7 viruses were accompanied by inefficient proteolytic processing of spike. B.1.1.7 viruses failed to escape from neutralizing antibodies, but slightly dampened induction of innate immunity. The bronchial cell line NCI-H1299 supported 24- and 595-fold increased growth of B.1.1.7 and B.1.617.2 viruses, respectively, in the absence of detectable ACE2 expression and in a spike-determined fashion. Superior spread in NCI-H1299 cells suggests that VOCs employ a distinct set of cellular cofactors that may be unavailable in standard cell lines.

## INTRODUCTION

Since its emergence, severe acute respiratory syndrome coronavirus 2 (SARS-CoV-2) has genetically diversified, giving rise to variants with altered phenotypic properties (Rambaut et al. 2020). In May 2021 the WHO announced a scheme for labeling SARS-CoV-2 lineages with evidence for increased transmissibility, severity, and escape from immunity (*https://www.who.int/en/activities/tracking-SARS-CoV-2-variants/*). The B.1.1.7 lineage was labeled Alpha as it was the first variant of concern (VOC) (O’Toole et al., 2021). First detected in the United Kingdom in September 2020 (*https://virological.org/t/preliminary-genomic-characterisation-of-an-emergent-sars-cov-2-lineage-in-the-uk-defined-by-a-novel-set-of-spike-mutations/563*), it shows a 50-100% higher reproduction number than previously circulating virus (Volz et al., 2021). Furthermore, the estimated hazard of death associated with B.1.1.7 is 61% higher than with pre-existing variants (Davies et al., 2021). B.1.617.2 was identified in India in October 2020 and was labelled VOC Delta in May 2021. In July 2021, the ECDC reported an increase of weekly COVID-19 cases observed in 20 European countries of 64.3%. Delta is the currently predominant circulating strain.

Genetic hallmarks of B.1.1.7 comprise a set of mutations resulting in nonsynonymous changes within the spike gene including deletion of amino acids 69, 70 and 144 in the amino-terminal domain, an N^501^Y exchange in the receptor-binding domain, an A^570^D exchange in subdomain 1, P^681^H and T^716^I exchanges in the proximity of the furin cleavage site and the S1 / S2 domain junction, as well as S^982^A and D^1118^H in the S2 domain. In addition to SNPs in ORF1ab and nucleoprotein, ORF8 in variant B.1.1.7. contains a premature stop codon. The functional role of ORF8 in SARS-CoV-2 is unclear. A variant with deleted ORF8 that circulated in Singapore during spring 2020 showed limited evidence for changes in *in-vitro* transcription profile (Gamage et al., 2020) and clinical outcome (Young et al., 2020).

To date, a functional correlate for the enhanced transmission and pathogenicity of B.1.1.7 and B.1.617.2 is missing. Studies of viral loads demonstrated that B.1.1.7- and B.1.617.2-infected individuals shed viral RNA of increased levels (Jones et al., 2021; Wang et al., 2021) and for prolonged time (Kissler et al., 2021b; Ong et al., 2021). Individual mutations in B.1.1.7 and B.1.617.2 spike have been investigated with regards to protein structure (Yang et al., 2021), *in vitro* ACE2-binding (Ramanathan et al., 2021), spike processing (Liu et al., 2021; Lubinski et al., 2021a) and stability (Motozono et al., 2021), as well as fitness (Motozono et al., 2021). In contrast to B.617.2 (Hoffmann et al., 2021a; Mlcochova et al., 2021; Planas et al., 2021), B.1.1.7 displays only modest, if any, alteration of sensitivity to neutralizing antibodies (Hoffmann et al., 2021b; Shen et al., 2021; Widera et al., 2021), suggesting a limited contribution of antibody-dependent immune escape to the observed phenotype of B.1.1.7. *In vitro* and *in vivo* replication of B.1.1.7 was found to differ depending on the model used. Some epithelial cell cultures and hamster models showed equal, slightly superior, or inferior replication for B.1.1.7 (Brown et al., 2021; Nuñez et al., 2021; Touret et al., 2021; Ulrich et al., 2021), while B.1.1.7 generally exhibited marginally superior replication in primates and ferrets (Rosenke et al., 2021; Ulrich et al., 2021). However, animal models may be limited in their capability to reflect adaptive processes that occur in a virus establishing endemicity in humans. Here, we studied the replication of B.1.1.7 viruses in different cell and organ models as well as dwarf hamsters, and identified a human cell line that reflects the replicative phenotype of B.1.1.7 as well as of B.1.617.2.

## RESULTS

### B.1 and B.1.1.7 SARS-CoV-2 display similar replication kinetics in immortalized cell lines

We first studied virus replication kinetics in a panel of immortalized cell lines. Compared to an early lineage (Non-VOC) B.1 strain carrying the D^614^G mutation (BavPat1/2020, (Wölfel et al., 2020)), two different clade B.1.1.7 isolates displayed smaller plaque size three days post-infection (Fig. 1A) and a delayed manifestation of cytopathogenic effects (CPE) in Vero E6 cells (Fig. 1B and Supplementary Movies), accompanied by a delay in infectious particle production during the initial 40 hours post-infection of Vero E6 cells (Fig. 1C).

**Figure 1.**
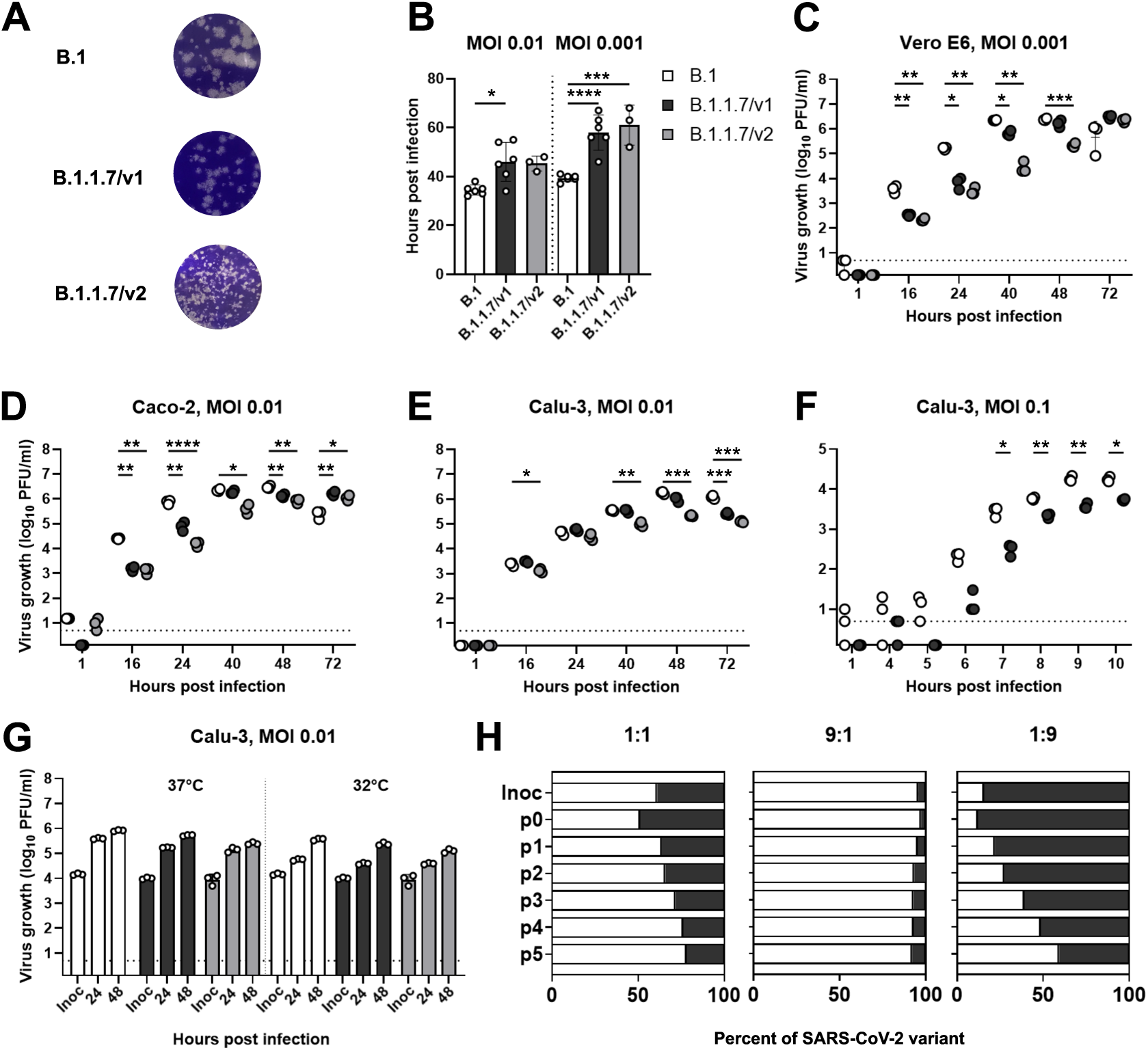
B.1 and B.1.1.7 SARS-CoV-2 display similar replication kinetics in immortalized cell lines. (A) Plaque morphology on Vero E6 cells which were infected with 1:100 diluted (B.1 and B.1.1.7/v1) or undiluted (B.1.1.7/v2) supernatants of infected Vero E6 cells. (B) Vero E6 cells were infected at the indicated MOI and onset of CPE was monitored by live cell imaging until 70 hours post-infection. (C-E) Virus growth was quantified in Vero E6 (C) Caco-2 (D) and Calu-3 (E) cells infected at indicated MOIs. Supernatant collected at the respective time points was titrated by plaque assay. Growth kinetic experiments in Vero E6 and Caco-2 cells were each performed in triplicates. One representative experiment out of two is shown for Calu-3 cells. (F) Virus growth in Calu-3 cells infected at an MOI of 0.1 was quantified at early time points after infection. (G) Virus growth kinetics in Calu-3 cells at 37°C (left) and 32°C (right). (H) Competition assay. Calu-3 cells were infected with a mixture of B.1 and B.1.1.7/v1 at indicated ratios. After serial passaging, viral RNA from the supernatant was isolated, sequenced and the relative proportion of B.1- and B.1.1.7-corresponding sequences (discriminated by a mutation in nsp12) was plotted. Data show arithmetic means of one experiment performed in triplicates. Dashed horizontal lines indicate the lower detection limit of the plaque assay. Inoc.: Inoculum, MOI: multiplicity of infection, PFU: plaque forming units, p0-p5: passage 0 - passage 5.

Unlike Vero E6 cells, Caco-2 and Calu-3 cells express the transmembrane protease serine subtype 2 (TMPRSS2) and are capable of producing type I interferons (IFNs). They supported the growth of B.1. equally or more efficiently than B.1.1.7 (Fig. 1D-E). Interestingly, B.1.1.7 production was particularly delayed in the very early phase of replication (Fig. 1F). Cultivation of infected cells at 32°C in order to resemble the temperature in the upper respiratory tract did not alter relative replication efficiencies (Fig. 1G). Under competitive passaging in Calu-3 cells, B.1 outcompeted B.1.1.7, even when the starting inoculum contained a nine-fold excess of B.1.1.7 (Fig. 1H). In sum, immortalized cell models failed to establish a clear growth advantage of B.1.1.7 that correlates with its enhanced transmissibility and pathogenicity *in vivo*.

### Absence of detectable fitness advantages of B.1.1.7 in primary human respiratory cells, organoids and hamsters

To detect potential variant-specific differences that may not become evident in immortalized cell lines, we studied more complex, physiologically relevant, primary human models of mucosal infection. Infection experiments in differentiated air-liquid interface cultures of human nasal (Fig. 2A), human bronchial (Fig. 2B) airway epithelial cultures (hBAECs), as well as epithelial intestinal organoids (Fig. 2C) failed to reveal a growth advantage for B.1.1.7 isolates compared to B.1. Growth of B.1.1.7 was even slightly inferior to B.1 in adult stem cell-derived human lung organoids (Fig. 2D). Virus production in the lung of infected dwarf hamsters was not different between any of the viruses used. In addition, virus production in the lungs of contact animals co-housed with infected animals did not show variant-specific quantitative differences in preliminary analyses (Fig. 2E).

**Figure 2.**
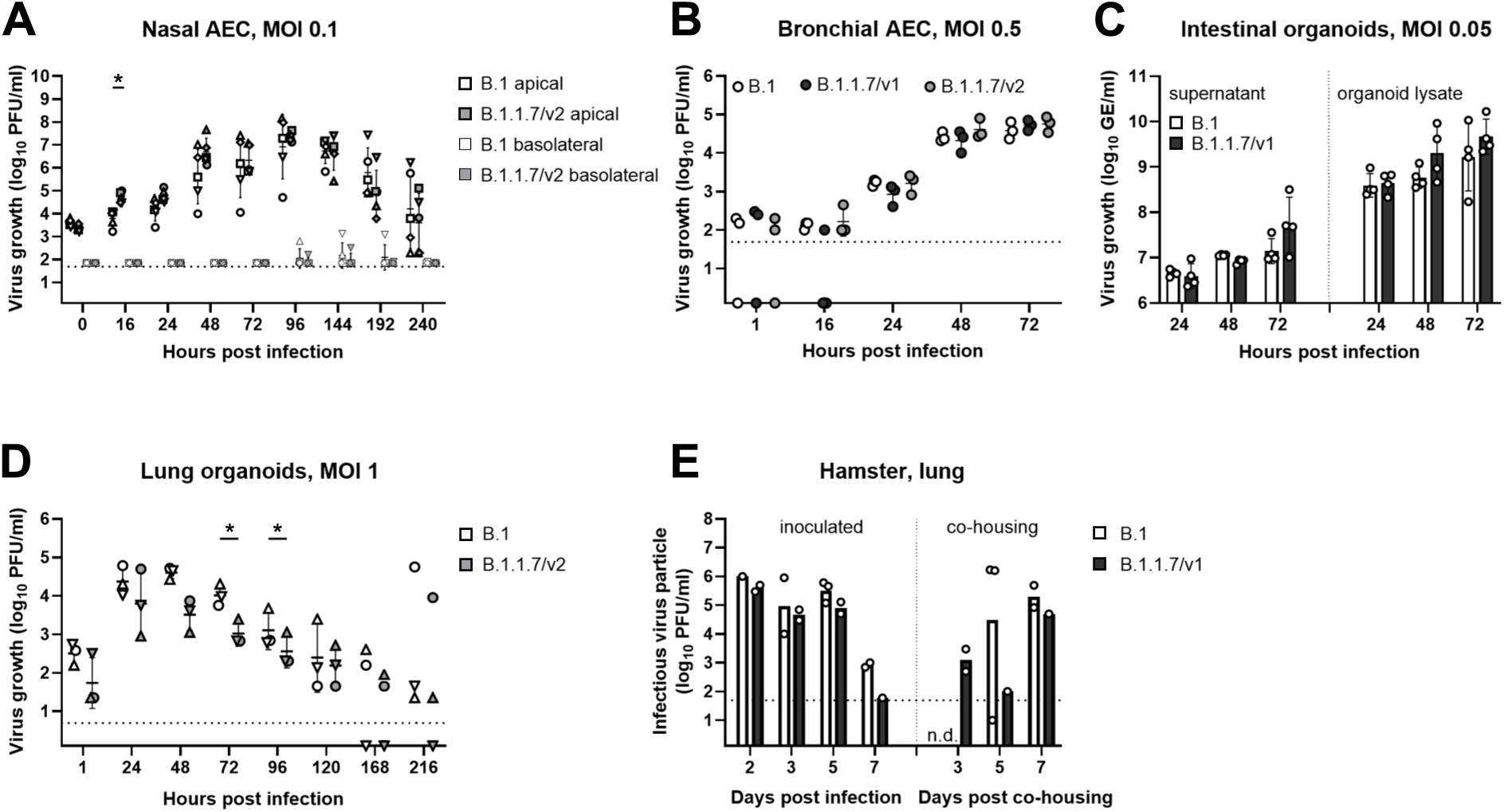
Comparison of fitness in primary human respiratory cells, organoids and dwarf hamsters. (A) Virus growth kinetics were performed in infected human nasal airway epithelial cultures (hNAECs) (MOI 0.1). Samples were collected from the apical and basal side at indicated time points and titrated by plaque assay. n=3 biological replicates. (B) Virus growth kinetics was conducted in infected bronchial AEC (MOI 0.5). Samples were collected from the apical side and titrated by plaque assay. Data are derived from one experiment conducted in triplicates. (C) Intestinal organoids were infected (MOI 0.05) and viral load in supernatant (left) and organoid lysates (right) was quantified at indicated time points by E-gene specific quantitative RT-PCR. Data are derived from four independent experiments. (D) Virus replication was monitored in infected lung organoids (MOI 1). Samples harvested at indicated time points were titrated by plaque assay. Data are derived from three independent experiments. (E) Dwarf hamsters were intranasally infected (100,000 PFU) and infectious virus particles from lung homogenates were quantified using plaque assay (left). Donor hamsters were co-housed with naive animals and transmission efficiency was determined from lung homogenates at the indicated time points (right). n=1-3 animals per experimental condition. Dotted horizontal lines indicate the lower detection limit of the plaque assays. GE: genome equivalents, n.d.: not detected

### B.1.1.7 spike protein shows decreased proteolytic processing

To identify potential consequences of B.1.1.7 spike mutations on expression and proteolytic processing of the glycoprotein, we analyzed lysates of HEK293T cells transfected with plasmids expressing SARS-CoV-2 spike-HA. Overall expression levels of spike constructs encoding individual or all B.1.1.7-specific mutations did not differ significantly from B.1 constructs in quantitative immunoblots (Fig. 3A, Fig. S1A). However, quantification of the proportion of S2-HA spike relative to the total spike-HA signal revealed a 1.8-fold reduction of proteolytic processing of the B.1.1.7 glycoprotein compared to that of B.1 (Fig. 3A). No single B.1.1.7-defining mutation, including P^681^H, fully recapitulated this property, even though deletion of H^69^/V^70^ showed a trend towards more efficient processing, in agreement with (Meng et al., 2021), and T^716^H showed a trend towards decreased proteolytic processing. The data suggest that a combination of amino acid exchanges is required for rendering proteolytic processing of B.1.1.7 spike less efficient. Reduced processing of B.1.1.7 spike was accompanied by a 2.3-fold decrease of spike levels associated with lentiviral particles, when compared to particles containing B.1 spike (Fig. 3B). Interestingly, the individual T^716^H mutation was sufficient for reduction of spike quantities associating to lentiviral particles. In accordance with previous reports (Kemp et al., 2021; Meng et al., 2021), deletion of H^69^/V^70^ increased spike abundance in pseudotyped particles.

**Figure 3.**
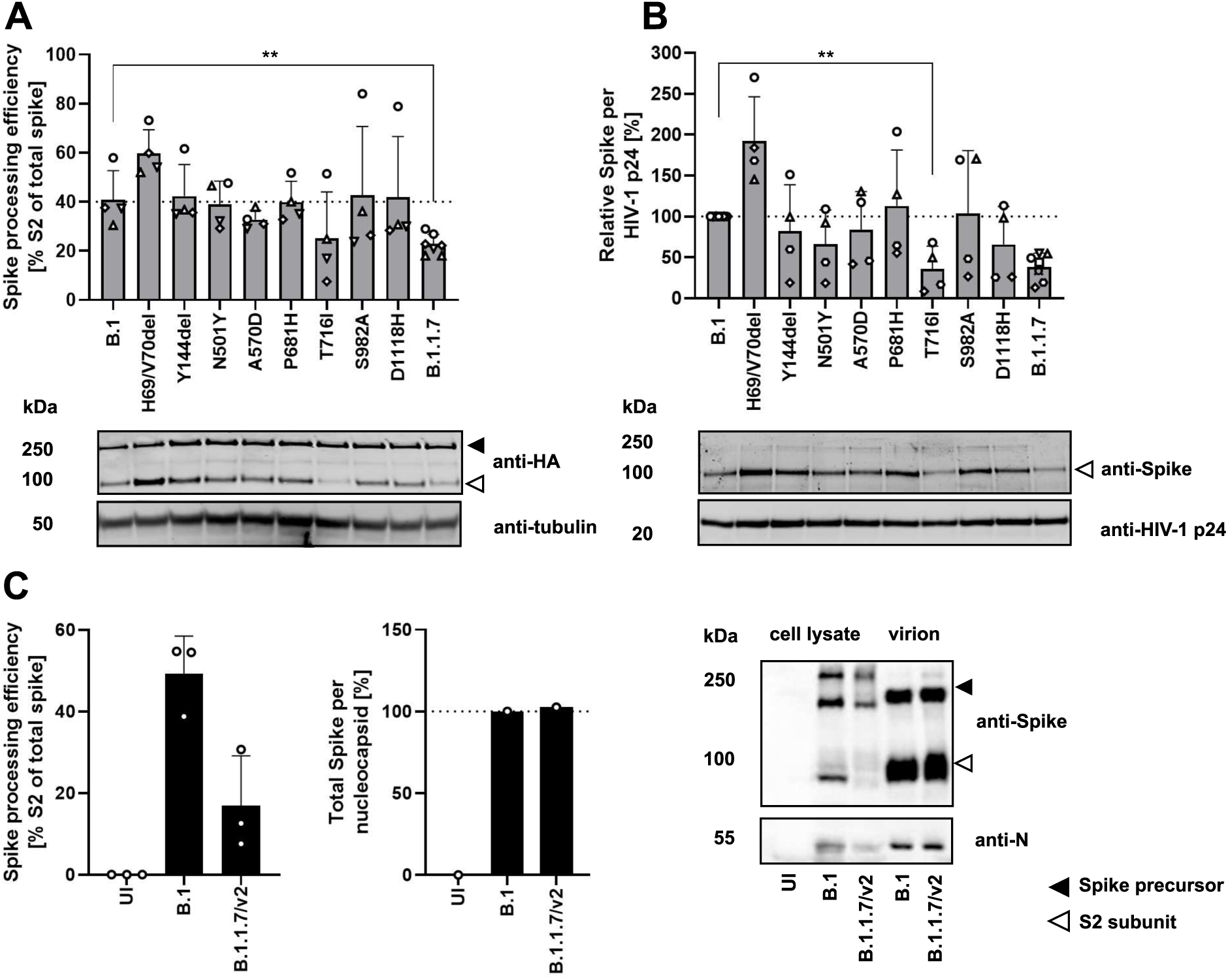
B.1.1.7 spike displays decreased proteolytic processing. (A) Spike processing in lysates of HEK293T cells expressing empty vector or SARS-CoV-2 spike-HA encoding individual or all B.1.1.7-corresponding mutations was quantified by immunoblotting (upper panel). Shown is one representative immunoblot (bottom panel) out of four. (B) Protein in lysed lentiviral particles pseudotyped with SARS-CoV-2 spike-HA was quantified by immunoblotting (upper panel). Shown is one representative immunoblot (bottom panel) out of four. (C) Vero E6 cells were infected with SARS-CoV-2 (MOI 5). Cells and virus-containing supernatants were harvested at 48 hours post-infection and processed for detection of spike and nucleocapsid by immunoblotting. Processing of spike in cell lysates (left panel) and spike incorporation in concentrated virion preparations (middle panel) was quantified. One representative blot out of two is shown (right panel). Black and white arrowheads indicate the bands of the uncleaved spike-HA precursor and of the cleaved S2-HA subunit, respectively. Statistical significance was calculated by a two-tailed, paired Student’s T-test. kDa: kilodalton, UI: uninfected

We challenged these findings by analysis of SARS-CoV-2-infected cells. B.1.1.7 spike in infected Vero E6 cells showed a 2.4-fold reduced processing efficiency (Fig. 3C, left panel and immunoblot, Fig. S1B). In virus particles, the ratio of spike per nucleocapsid signal appeared intact, suggesting that inefficient proteolytic cleavage does not translate into a decreased association of mature S2 into virions (Fig. 3C, middle panel and immunoblot). Overall, expression levels of spike did not differ significantly from B.1 in quantitative immunoblots (Fig. S1C).

### Enhanced cell-cell fusion and reduced virus particle entry by B.1.1.7 SARS-CoV-2 spike

We next analyzed fusogenicity of individual spike proteins in a cell-cell fusion assay based on co-cultures of CHO cells transiently expressing HIV-1 Tat and individual SARS-CoV-2 spike proteins, and ACE2/TMPRSS2-transfected, LTR-luciferase-expressing target TZM-bl cells. Compared to B.1 spike, no significant changes in membrane fusion activity were detected with any single mutation present in B.1.1.7 spike (Fig. 4A). However, full B.1.1.7 spike was more prone to induce cell-cell fusion. Furthermore, entry of lentiviral pseudotypes mediated by the same set of spike proteins was quantified after transduction of Calu-3 cells using p24 capsid normalized inocula in a luciferase-based assay (Fig. 4B). Whereas most individual mutations did not significantly alter the ability of SARS-CoV-2 spike to mediate entry into Calu-3 cells, T^716^I and the full B.1.1.7 spike mediated reduced entry (Fig. 4B, Supplemental Fig. S2A). The inferior ability of B.1.1.7 spike-containing lentiviral pseudotypes to transduce Calu-3 cells was corroborated in titration experiments on ACE2/TMPRSS2-positive A549 cells (Fig. 4C) and is potentially related to the lower levels of incorporated spike when compared to B.1 spike-pseudotyped particles.

**Figure 4.**
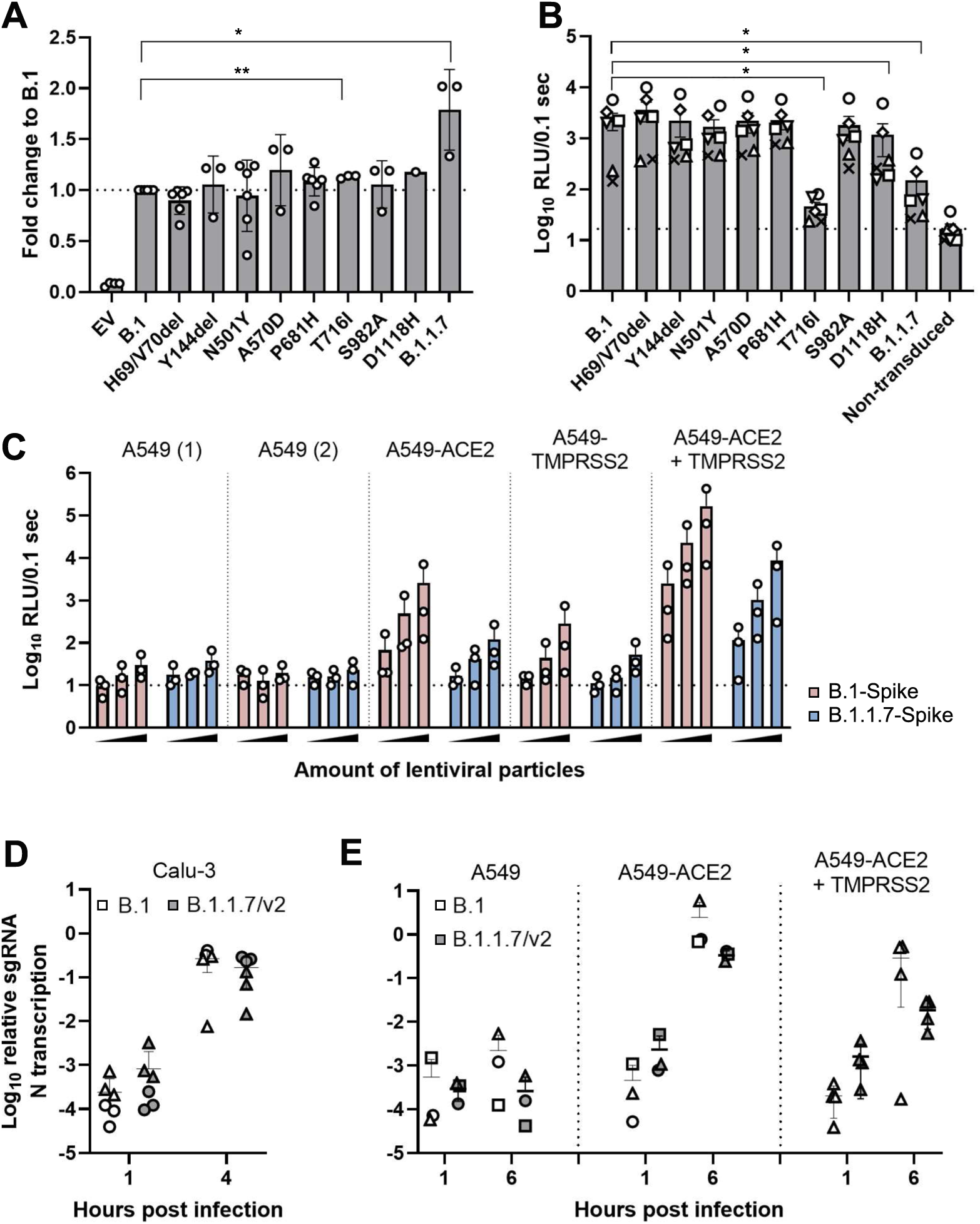
Membrane fusion and entry based on lentiviral pseudotypes and SARS-CoV-2 particles. (A) For Tat-mediated cell-cell fusion assay, CHO cells were co-transfected with plasmids expressing indicated spike-HA and HIV-1 Tat. LTR-luciferase-expressing target TZM-bl cells were transfected with plasmids encoding human ACE2 and TMPRSS2. Transfected cells were co-cultured for eight hours and luciferase expression resulting from intercellular Tat transfer was quantified luminometrically. All values were normalized to B.1 spike (indicated by a dotted line). Shown are results from three-six biological replicates, each performed in triplicates. (B) Calu-3 cells were transduced with lentiviral pseudoparticles expressing luciferase and decorated with indicated spike-HA. Transduction efficiency was quantified luminometrically. Dotted line indicates background levels of luciferase non-transduced cultures. Shown are results from six biological replicates, each performed in triplicates, indicated by symbols. (C) Indicated A549 cells were transduced with increasing quantities (0.5 µl, 5 µl and 50 µl) of lentiviral, luciferase-expressing particles pseudotyped with B.1- or B.1.1.7-spike. Transduction efficiency was determined luminometrically. Dotted line indicates luciferase background level of luciferase detected in non-transduced cells. Symbols represent individual values of three biological replicates, each performed in triplicates. (D, E) Calu-3 (D) and indicated A549 (E) cells were infected at 4°C with B.1 or B.1.1.7 isolates (MOI 1) to allow synchronized infection. Total cellular RNA was isolated at the indicated time points and nucleocapsid-encoding subgenomic RNA was quantified by RT-PCR. Del: deletion, RLU: relative light units, sgRNA N: subgenomic nucleocapsid RNA, conc.: concentration

We next investigated entry kinetics in Calu-3 and A549-ACE2 cells with authentic SARS-CoV-2. In single-cycle entry experiments, virus inocula were absorbed at 4°C for one hour, cells were washed to remove excessive virus particles, and the cells eventually incubated at 37°C to initiate synchronized entry. *De novo*-synthesized, cell-associated subgenomic transcripts served for detection of early signs of virus replication after entry (Fig. 4D and 4E). Reminiscent of our results obtained with the lentiviral pseudotypes, B.1.1.7 initiated slightly lower levels of virus replication in Calu-3, A549-ACE2 and A549-ACE2/TMPRSS2 cells when compared to B.1 at the earliest time point at which subgenomic RNA production was detectable. ACE2 seems to be required for entry into A549, as levels of subgenomic viral RNA remained at background levels in parental A549. Entry of both viruses in Calu-3 cells depended on TMPRSS2 and furin and less on a cathepsin L-dependent pathway. In addition, clathrin inhibition resulted in decreased endocytic entry of B.1 and B.1.1.7 in Calu-3 cells (Fig. S2B), as was previously described for SARS-CoV (Wang et al., 2008). Inocula were back-titrated to ensure that equal amounts of virus were used for infection. In conclusion, B.1.1.7 spike-pseudotyped lentiviral particles and authentic B.1.1.7 are less efficient in entering susceptible target cells than their B.1 counterparts. This inefficiency may be related to the detected lower efficiency of spike processing.

### B.1.1.7 fails to escape from neutralizing antibodies and may dampen induction of innate immunity

In accordance with reports by others (Bates et al., 2021; Supasa et al., 2021; Widera et al., 2021), B.1 infection- and vaccination-induced antibodies efficiently neutralized both B.1 and B.1.1.7 in a plaque reduction neutralization test, whereas such antibodies failed to effectively neutralize B.1.351 (Fig. 5A). Interestingly, binding of B.1.1.7 RBD, which differs from that of B.1 solely at one position (N^501^Y), to ACE2 in a surrogate neutralization test was slightly less sensitive to inhibition by antibodies from non-VOC convalescents and vaccinees when compared to antibodies raised following B.1.1.7 infection (Fig. 5B), indicating that antibodies targeting other regions than the RBD may contribute to the neutralization of infection. However, B.1.351 RBD was clearly more resistant to ACE2 binding inhibition than the B.1 and B.1.1.7 RBDs. These results suggest a large absence of escape from humoral immunity by B.1.1.7, as opposed to B.1.351.

**Figure 5.**
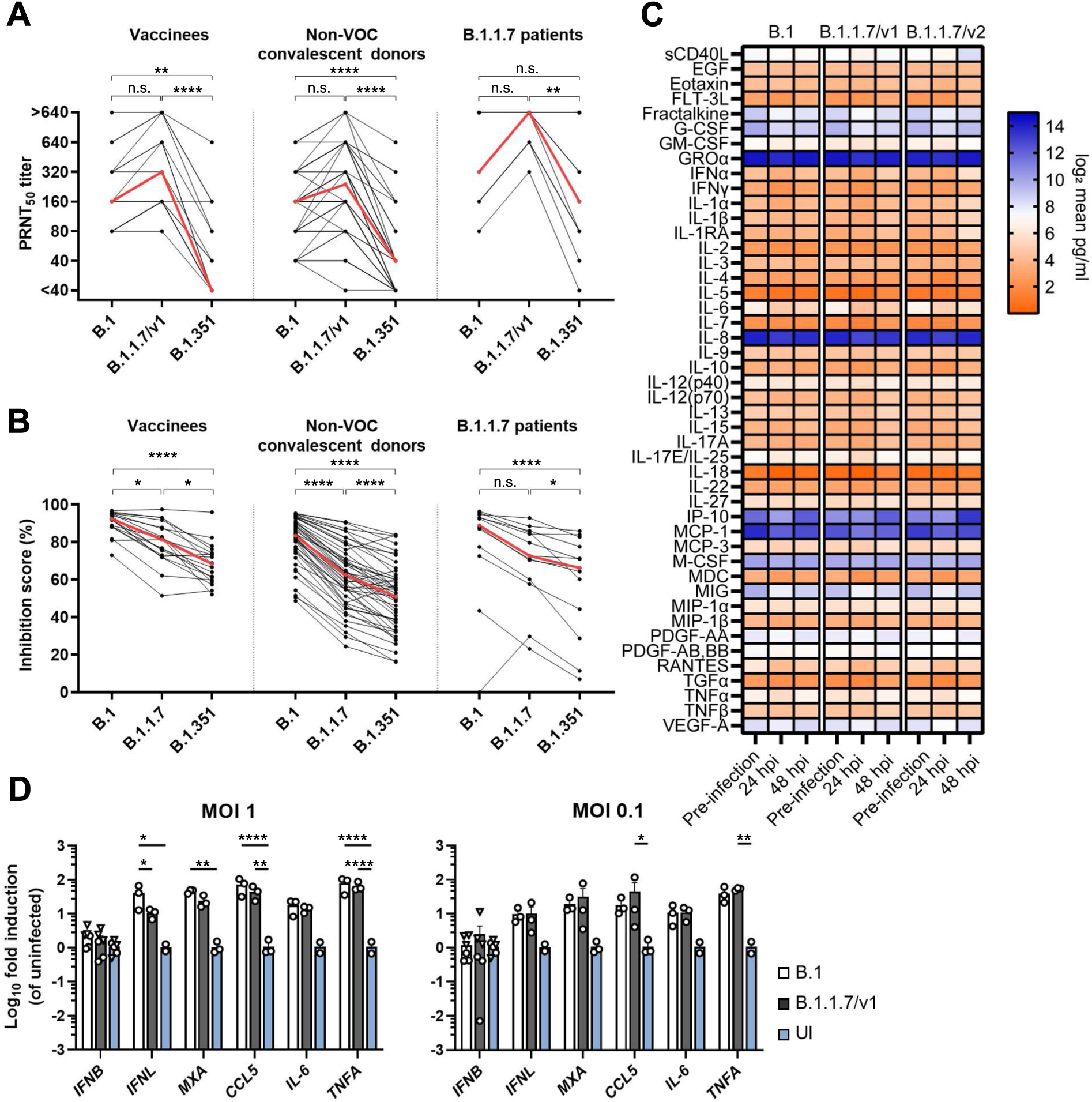
B.1.1.7 fails to escape from neutralizing antibodies and may dampen induction of innate immunity. (A) Neutralizing titers against the indicated virus strains were determined in plaque reduction neutralization tests (PRNT). Red line indicates median titers per group. (B) Inhibition of ACE2/RBD interaction was measured using surrogate virus neutralization assays. Sera were tested using RBD proteins of B.1, B.1.1.7 and B.1.351 as indicated. Red lines indicate median values. The same set of samples was measured in (A) and (B), vaccinees n=19, Non-VOC convalescent donors n=50, B1.1.7 patients n=13. (C) Concentration (pg/ml) of cytokines and chemokines in the basal medium of infected bronchial airway epithelial cells (MOI 0.5). Concentration of cytokines and chemokines was determined by MagPix Luminex technology. (D) Calu-3 cells were infected at indicated MOIs or left uninfected (UI) and cell lysates were generated 16 hours post-infection. Total RNA was extracted and expression of the indicated genes was determined by quantitative real-time PCR. Shown is the mean fold change +/− S.D.. The experiment was performed in triplicates.

We next hypothesized that B.1.1.7 has evolved superior ability to prevent or evade cell-intrinsic immunity. In the basal medium of infected differentiated bronchial airway epithelial cell cultures, no significant variant-specific differences were identified for various cytokines and other secreted proteins related to innate immunity, including IFN-α, IFN-γ, and IP-10 (Fig. 5C). In infected Calu-3 cells, expression of *IFNB*, *MXA*, *CCL5*, *IL6* and *TNFA* was induced to similar levels by both variants (Fig. 5D). Interestingly, B.1.1.7 infection appeared to induce slightly lower levels of *IFNL* expression than B.1 (Fig. 5D), suggesting that B.1.1.7 may be able to suppress innate immune responses more efficiently.

In conclusion, absence of a detectable escape from neutralizing antibodies suggest that the enhanced B.1.1.7 transmissibility and pathogenicity *in vivo* involves other immune evasion mechanisms, potentially the ability to dampen induction of an antiviral state in infected cells.

### B.1.1.7 and B.1.617.2 display a spike-dependent, ACE2-independent post-entry replication advantage in NCI-H1299 cells

We reasoned that common cell culture models used up to this point may have failed to reflect the predominant spread of B.1.1.7 observed in the human population. Upon further exploration of susceptible cell lines, we identified that the human bronchial cell line NCI-H1299 (Phelps et al., 1996) supports B.1.1.7 growth with up to 24-fold higher efficiency than B.1 growth (Fig. 6A) in multi-cycle growth kinetics. The smaller plaque size of B.1.1.7 virus originating from Vero E6 cells (see Fig. 1A, lower panel) was corroborated with NCI-H1299 cell-derived virus.

**Figure 6.**
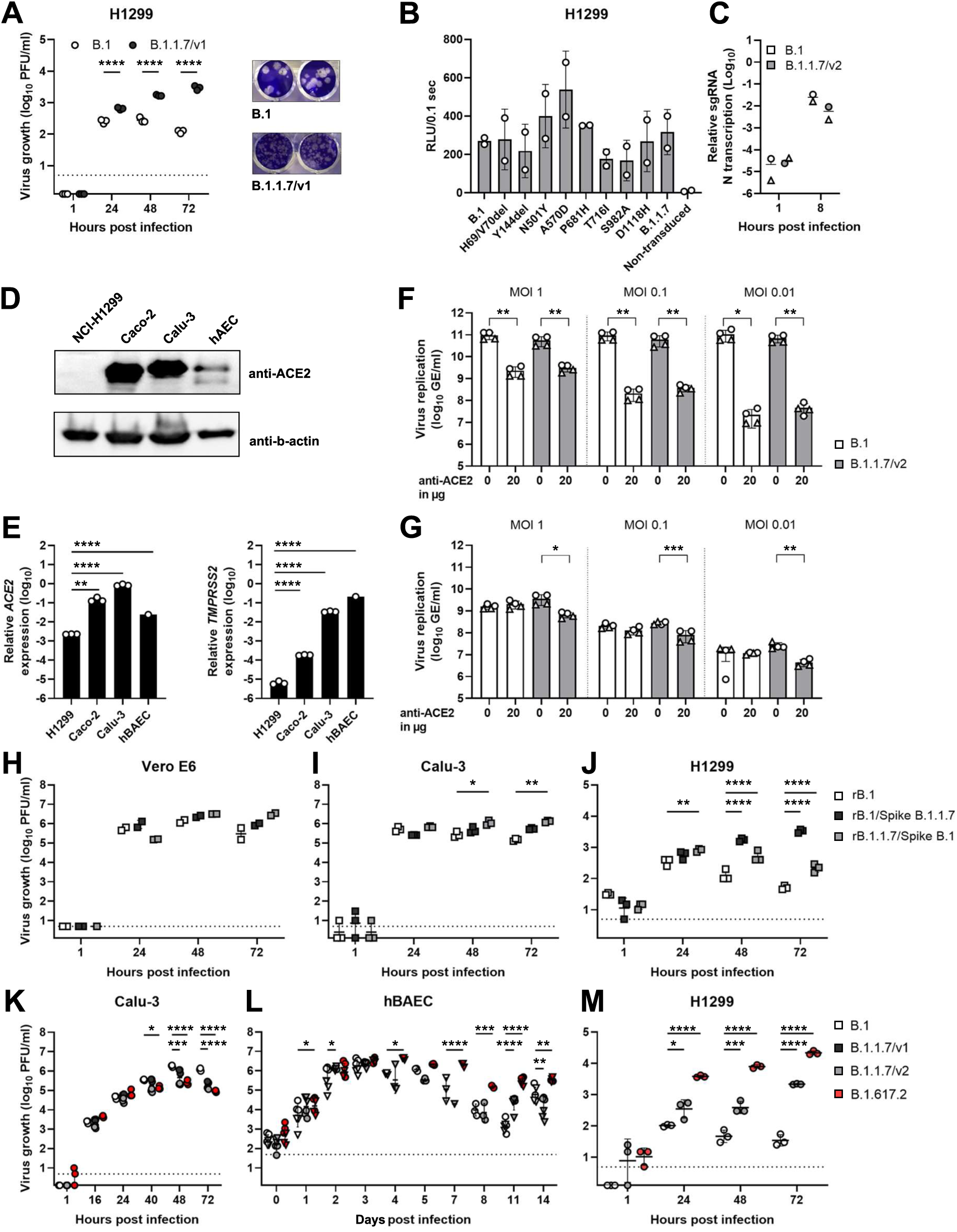
B.1.1.7 and B.1.617.2 display a spike-dependent, ACE2-independent post-entry replication advantage in NCI-H1299 cells. (A) Virus growth of B.1 and B.1.1.7/v1 was assessed on NCI-H1299 cells. Cells were infected (MOI 0.01) and supernatants of indicated time points were titrated on Vero E6 cells. Plaque morphology of NCI-H1299-derived B.1 and B.1.1.7/v1 on Vero E6 cells is shown. (B) NCI-H1299 cells were transduced with increasing amounts of lentiviral particles pseudotyped with indicated spike proteins. Pseudotype entry was analyzed luminometrically in cell lysates. Data from two biological replicates, each performed in triplicates, is shown. White symbols represent arithmetic means of the biological replicates. (C) H1299 cells were infected in triplicates at 4°C with B.1 or B.1.1.7 isolates (MOI 1) to allow synchronized entry. Relative quantities of cell-associated nucleocapsid-specific subgenomic RNA were determined by quantitative RT-PCR. Two independent experiments were performed, each conducted in 3-4 replicates. Symbols represent the arithmetic means of each experiment. (D) ACE2 expression levels were analyzed by immunoblotting. Beta-actin was used as a loading control. (E) Expression of *TMPRSS2* and *ACE2* was quantified by quantitative RT-PCR in indicated cells. (F, G) Calu-3 (F) and NCI-H1299 (G) cells were pretreated with 20 µg/ml anti-ACE2 antibody for one hour prior to infection with B.1 and B.1.1.7 isolates (MOI of 0.01). At 48 hours post-infection, viral replication was quantified from the supernatant by the use of E-gene assay. Results from two independently performed experiments, each conducted in triplicates, are shown. (H-J) Vero E6 (H), Calu-3 (J) and NCI-H1299 (J) cells were infected (MOI 0.01) and supernatant was titrated on Vero E6 cells. The growth experiment in Vero E6 cells was performed once in duplicates. Growth experiments in Calu-3 and NCI-H1299 cells were performed once in triplicates. (K-M) Virus growth of B.1, B.1.1.7 and B.1.617.2 isolates (MOI 0.01) was quantified in Calu-3 (K), human bronchial airway epithelial cells (hBAECs) (L) and NCI-H1299 (M) cells. Dashed horizontal lines indicate the lower limit of detection of the plaque assay.

Interestingly, there was no advantage in entry of lentiviral particles based on the B.1.1.7 spike protein in NCI-H1299 cells (Fig. 6B). Also, there was no advantage for authentic SARS-CoV-2 B.1.1.7. virus in entering NCI-H1299 cells in a synchronized entry assay (Fig. 6C), indicating that the more efficient replication of B.1.1.7 may not necessarily be determined by improved entry. Even more surprisingly, replication of B.1.1.7 in NCI-H1299 cells occurred in the absence of detectable ACE2 protein (Fig. 6D), and *ACE2* and *TMPRSS2* mRNAs were only weakly expressed as compared to other SARS-CoV-2-susceptible cell cultures (Fig. 6E). In order to exclude the possibility that SARS-CoV-2 infection in NCI-H1299 cells was maintained by minute traces of ACE2 expressed below the detection limit of our system, we blocked ACE2 by antibodies. Individual incubation of Vero E6 and Calu-3 cells with three ACE2-neutralizing antibodies abolished and diminished SARS-CoV-2 infection, respectively (Fig. 6F, Fig. S3A-B). In contrast, anti-ACE2 antibody treatment did not and only very modestly modulate the susceptibility of NCI-H1299 to B.1 and B.1.1.7 SARS-CoV-2 infection (Fig. 5G and Fig. S3C), reinforcing the notion that this cell line supports infection via a largely ACE2-independent mechanism.

To analyze the specificity of the observed phenotypic changes for the spike protein, we generated, by reverse genetics, a Wuhan-1 virus with a D^614^G mutation (representing a B.1 virus, rB.1), as well as a Wuhan-1 virus carrying the full spike protein of B.1.1.7 (rB.1/Spike B.1.1.7). Moreover, we engineered a virus consisting of the B.1.1.7 backbone but the Wuhan-1 D^614^G spike, representing a prototypic B.1 spike (rB.1.1.7/Spike B.1, Fig. 5H-J). As expected, all recombinant viruses grew to similar titers in Vero E6 cells (Fig. 6H). In Calu-3 cells, growth of rB.1.1.7/Spike B.1 was increased 9-fold over rB.1 at 72 hours post-infection, suggesting that the B.1.1.7 backbone may confer replication advantages that do not depend on the spike protein (Fig. 6I). In NCI-H1299 cells, the virus expressing the B.1.1.7 spike in the backbone of B.1 grew to considerably higher titers (up to 65-fold increased titer) than the reciprocal virus and the original rB.1 virus (up to 15-fold increased titer, Fig. 6J). This suggests the replicative advantage of B.1.1.7 in NCI-H1299 cells, while probably exerted on entry, is mediated by the spike protein.

We next reasoned that NCI-H1299 cell-specific increased viral growth may be a property that is shared by the current, predominating VOC Delta. In Calu-3 cells and in hBAEC cultures, a B.1.617.2 isolate grew to similar and increased titers, respectively, compared to B.1 and B.1.1.7 (Fig. 6K and Fig. 6L). Remarkably, in multi-cycle infection experiments in NCI-H1299 cells, B.1.617.2 grew to 264 (min. 37 -max. 595)-fold higher titers than B.1 and 24 (min. 3 -max. 59)-fold higher titers than B.1.1.7, respectively (Figure 6M). Together, these data suggest that VOCs B.1.1.7 and B.1.671.2 may utilize partially common mechanisms of replication enhancement that are spike-dependent but do not improve entry efficiency.

## DISCUSSION

To date, our understanding of molecular mechanisms that underlie the rapid spread and/or increased pathogenicity of SARS-CoV-2 VOCs is still in its infancy. As SARS-CoV-2 productively infects epithelial cells (Hou et al., 2020), and B.1.1.7-infected patients shed 10-fold more viral RNA (Jones et al., 2021), we hypothesized that B.1.1.7 has a replication advantage in human epithelial cell cultures. Although all immortalized, primary, and organoid cultures tested were highly permissive for SARS-CoV-2 infection, we failed to detect a B.1.1.7-specific growth advantage with one notable exception. Our findings corroborate reports by others showing a similar growth rate of B.1.1.7 and B.1. viruses in common culture cells and primary human airway epithelial (HAE) cells (Brown et al., 2021; Touret et al., 2021; Ulrich et al., 2021). Despite monitoring infected cultures for up to ten days, we failed to observe temporally increased virus production in human organoid models as seen by others (Lamers et al., 2021). *In vivo*, B.1.1.7 has been proposed to display a fitness advantage in experimentally infected ferrets, in hACE2-K18Tg-transgenic mice and hACE2-KI transgenic mice (Ulrich et al., 2021), but not in Syrian hamsters (Nuñez et al., 2021; Ulrich et al., 2021). In line with these results, our data confirm the absence of significant growth differences between B.1.1.7 and B.1 variants in experimentally infected dwarf hamsters (Abdelnabi et al., 2021). However, a recent study did observe a transmission advantage of B.1.1.7 upon low dose infection of Syrian hamsters (Mok et al., 2021). Finally, in intranasally infected African green monkeys, B.1.1.7 infection generated higher levels of viral RNA in the respiratory tract than B.1 infection (Rosenke et al., 2021). Together, the superiority of B.1.1.7 spread can be recapitulated in some animal models, but is not detectable in most cell culture systems.

B.1 SARS-CoV-2 productively infects susceptible cells via binding of the spike protein to ACE2 and TMPRSS2-mediated priming of spike (Hoffmann et al., Cell 2020). Compared to B.1 viruses, B.1.1.7 viruses and B.1.1.7 spike-decorated lentiviral particles were equally or even less efficient in entering ACE2-expressing cell lines. Intriguingly, B.1.1.7 spike appeared to be more fusogenic in cell-cell fusion assays, in accordance with published work (Meng et al., 2021; Rajah et al., 2021). Proteolytic processing and viral packaging of coronaviral spike can be rate-limiting during infectious virus production. Our findings of reduced proteolytic cleavage in spike plasmid-transfected and SARS-CoV-2-infected cells are consistent with reduced furin-mediated processing of B.1.1.7 spike as shown by a biochemical peptide cleavage assay (Lubinski et al., 2021b), and challenge data reporting intact processing of cell-associated spike when expressed in spike plasmid-transfected cells (Mlcochova et al., 2021). Of note, the latter study used plasmids expressing spikes lacking 19 C-terminal amino acids, an experimental modification that has been widely accepted for artificial enhancement of cell surface expression and lentiviral incorporation of spike (Yu et al., 2021), and that we refrained to adopt in order to maintain the expression context as physiological as possible. The observation of reduced processing translated into lower levels of lentivirus-associated spike, which may be the cause of the reduced infectivity of particles. Even though impaired maturation did not detectably alter virion-associated spike levels under our experimental conditions, we cannot exclude that it may still modulate the kinetics of virus particle secretion and/or the quality of secreted SARS-CoV-2 particles.

Whereas increased B.1.1.7 replication has been observed in patients, the initial phase of virus infection is difficult to capture in clinical observations due to late sampling, making cell culture studies potentially more insightful. On the contrary, cell cultures may fail to reflect differences in virus production in later stages of tissue infection due to the limiting effect of cytopathogenic effects i*n vitro*. Our observation of reduced growth of B.1.1.7 in cell culture does not necessarily contradict clinical observations. We observed a slower ramp-up of virus production, delayed onset of cytopathic effect, and slightly reduced levels of *IFNL* expression for B.1.1.7, which, overall, is consistent with a stealthy invasion of tissue with initially limited production of PAMPs and more efficient evasion of cell-intrinsic innate immunity, as suggested by others. The previously reported transcriptional changes affecting the N gene reading frame, leading to expression of ORF9b potentially acting as an antagonist of IFN induction, as well as the reported increase of production of the IFN signaling antagonist ORF6, may contribute to this phenotype (Parker et al., 2021; Thorne et al., 2021). Accordingly, infected air-liquid interface cultures of alveolar type 2 cells produced more infectious B.1.1.7 virus when compared to B.1 only in a late phase of infection (Lamers et al., 2021). Also, replication of B.1 and B.1.1.7 variants in the upper respiratory tract of experimentally infected African green monkeys differed only from day 5 post-infection (Rosenke et al., 2021). Together, this suggests a delayed but extended phase of infectious B.1.1.7 virion shedding, potentially resulting in a prolonged phase of heightened transmission probability, which is also supported by preliminary viral load studies (Jones et al., 2021; Kissler et al., 2021a).

While cell-intrinsic immunity may be involved, its onset and extent is still dependent on fundamental factors such as receptor usage and cell entry, often reflected by changes in the spike protein. Paradoxically, ACE2 expression levels in the respiratory tract are low and common cell culture models may not fully recapitulate *in vivo* properties of primary target tissues of SARS-CoV-2 infection (Hikmet et al., 2020). Our exploration of other cell cultures may thus have uncovered a cell line that has remained understudied with regard to SARS-CoV-2 infection phenotypes. While caveats and limitations of neoplastic cell lines apply, NCI-H1299 cells yielded higher levels of replication and infectious virus production not only for B.1.1.7, but also for B.1.617.2. This cell line of epithelial morphology is devoid of detectable ACE2 protein and remains susceptible to SARS-CoV-2 infection even in the presence of ACE2-neutralizing antibodies, suggesting the existence of an alternative, but not variant-specific mode of entry. This finding is reminiscent of a recent study (Puray-Chavez et al., 2021) that identified SARS-CoV-2 infection of the H522 lung cell line in an ACE2-independent fashion (in this study the alternative way of infection could only be utilized by viruses carrying an E^484^D substitution within the spike protein RBD).

Surprisingly, while experiments with reciprocal chimeras established that the B.1.1.7 spread advantage in NCI-H1299 cells is mediated to a large extent by its spike, the entry process *per se* was similarly or even less efficient for B.1.1.7 than for B.1. Notably, a virus consisting of the B.1.1.7 backbone with B.1 spike displayed a slightly better replication efficiency than the original B.1 virus in type-I IFN-competent Calu-3 cells, confirming that other genetic determinants of B.1.1.7 beyond spike contribute to enhanced replication. These findings suggest a post-entry, yet largely spike-dependent replication advantage of B.1.1.7 that manifests itself in NCI-H1299 cells. Remarkably, we conducted preliminary studies of replication for B.1.617.2 and found a highly similar phenotype, i.e., a similar level of replication in common ACE2-positive cell lines but a specific increase of replication in NCI-H1299 cells that does not seem to be determined by improved entry. In addition, B.1.617.2 displayed a spreading advantage in human BAECs that manifested itself not before four days post-infection. The extent of growth advantage for B.1.617.2 in NCI-H1299 cells was higher than for B.1.1.7, which would be expected given the observed differences in epidemic growth rates for these VOCs. More work will be necessary to understand the nature of the alternative entry mechanism, the identity of the entry-independent and variant-specific difference in replication as a function of spike, as well as the possibility that two VOCs may have undergone convergent evolution towards utilization of this same unknown mechanism. Also, whether the ACE2-independent entry is linked to the replication advantage, e.g. by potentially more efficient virion release due to ACE2 scarcity or absence, remains unclear at this time. Mere absence of ACE2 expression is not sufficient for B.1.1.7-specific growth advantage as illustrated by the refractoriness of ACE2-negative A549 cells to SARS-CoV-2 infection. Future studies are required for the identification of a hypothetical alternative receptor of SARS-CoV-2 and for the elucidation of the specific cellular environment of NCI-H1299 cells that confers a replication advantage to B.1.1.7 and B.1.617.2 SARS-CoV-2.

## MATERIALS AND METHODS

### Samples from COVID-19 patients and vaccinees

Sera were available through a study on convalescent plasma donors, who recovered from mild to moderate COVID-19 before the emergence of any SARS-CoV-2 VOCs (Schlickeiser et al., 2020). Additional sera were available through a study on SARS-CoV-2 infection and COVID-19 vaccination (Hillus et al., 2021), a prospective observational cohort study Pa-COVID-19 (Thibeault et al., 2021) including its study arm RECAST (Understanding the increased resilience of children compared to adults in SARS-CoV-2 infection) and from RT-PCR confirmed B.1.1.7-infected patients. The use of clinical samples (sera) was approved by the Institutional Review Board at Charité -Universitätsmedizin Berlin (EA1/068/20, EA2/092/20 and EA2/066/20) and is in accordance with the Berlin State Hospital Law, allowing for pseudonymized scientific analysis of routine patient data.

### Cells and culture conditions

A549 parental (ATCC CCL-185), A549-ACE2, A549-TMPRSS2, A549-ACE2 + TMPRSS2 (Widera et al. 2021), Caco-2 (ATCC HTB-37), Calu-3 (HTB-55), CHO (HIV Reagent Program ARP-2238), NCI-H1299 (ATCC CRL-5803), HEK-293T (ATCC CRL-3216), TZM-bl (HIV Reagent Program ARP-8129) and Vero E6 (ATCC CRL-1586) cells were maintained at 37°C and 5% CO_2_ in a humidified atmosphere and cultured in Dulbecco’s Modified Eagle’s Medium (DMEM, ThermoFisher Scientific) supplemented with 10% fetal bovine serum (FBS, ThermoFisher Scientific), 1% non-essential amino acids 100x concentrate (NEAA, ThermoFisher Scientific) and 1% sodium pyruvate 100 mM (NaP, ThermoFisher Scientific) and split twice a week. For seeding and cultivation, cells were washed with phosphate buffered saline (PBS, ThermoFisher Scientific) and detached with 0.05% trypsin-EDTA solution (ThermoFisher Scientific).

### Virus strains

Infection experiments were done with BetaCoV/Munich/ChVir984/2020 (B.1, EPI_ISL_406862), hCoV-19/Germany/BY-ChVir21652/2020 (B.1.1.7/v1, EPI_ISL_802995), BetaCoV/Baden-Wuerttemberg/ChVir21528/2021 (B.1.1.7/v2, EPI_ISL_754174) and hCoV-19/Germany/BW-ChVir22131/2021 (B.1.351, EPI_ISL_862149). A virus of the B.1.617.2 (“Delta”) clade (hCoV-19/Germany/SH-ChVir25702_4/2021) was isolated from a patient in Schleswig-Holstein, Germany, and its sequence deposited in Gisaid (EPI_ISL_2500366). Due to the observation of rapid cell culture-induced mutations at the spike polybasic furin cleavage motif (spike amino acid no. 681-685), all virus stocks were sequenced by next generation sequencing to confirm the absence of minority variants. Unless otherwise stated, only virus stocks with no or variant frequencies below 20% of all sequence reads were included in downstream infection experiments. Virus isolation and all SARS-CoV-2-related infection experiments were performed under biosafety level-3 (BSL-3) conditions with enhanced respiratory personal protection equipment.

### Plasmids

Codon-optimized, C-terminally tagged spike cDNAs in pCG were generated using pCG-SARS-CoV-2 spike Wuhan as a template (Hoffmann et al., 2020) in which the N-terminus was repaired and D^614^G was introduced via site-directed mutagenesis. The individual B.1.1.7-characteristic mutations were introduced individually and in combination by site-directed mutagenesis. All constructs were confirmed by Sanger sequencing.

### Virus isolation

Virus was isolated from naso- or oropharyngeal swabs using Vero E6 cells. Cells were seeded at a density of 175,000 cells per well in 24-well plates one day prior to isolation. For virus isolation, the medium was removed and cells were rinsed once with 1x PBS (ThermoFisher Scientific) and inoculated with 200 µl of swab sample. After one hour incubation, 800 µl of isolation medium (DMEM, supplemented with 2% FBS, 1% penicillin-streptomycin and 1% amphotericin B, ThermoFisher Scientific) was added to each well. Cells were monitored for cytopathic effects (CPE) every day. Four days post-inoculation, viral RNA was isolated and quantified from the supernatant as described below. Isolation success was determined when CPE was detectable and viral RNA concentrations were above a threshold of 100,000 genome equivalents per µl. Virus stocks were produced from all positive cultures.

### Virus stock production

Vero E6 cells were seeded in T175 tissue culture flasks alowing the cells to reach 90% confluence on the following day. Cells were washed once with PBS and inoculated with 100 µl of low passage (passage 1-2) virus stock solution (approximately 1,000,000 PFU per ml) in 20 ml virus infection medium (DMEM supplemented with 2% FBS, 1% NEAA, 1% sodium phosphate). Three days post-inoculation, supernatant was harvested and virus particles were purified from cytokines and concentrated using Vivaspin 20 (Sartorius, filtration units with a size exclusion of 100 kDa) according to the manufacturer’s instructions. Virus concentrate was resuspended in 2-3 ml PBS, diluted 1:2 in virus preservation medium (0.5% gelatine in OptiPRO serum free medium) and stored at −80°C. Infectious titers were determined in three independent plaque titration experiments and viral RNA concentration was quantified by real-time RT-PCR (E gene assay). All stocks were sequenced by next generation sequencing methods and the absence of additional mutations was confirmed to occur in less than 20% of the virus-specific reads.

### Virus infection and virus growth kinetics in cell cultures

Vero E6, Caco-2, NCI-H1299 and A549 cells were seeded at a densitiy of 350,000 cells per ml and Calu-3 cells at a density of 500,000 cells per ml one day prior to infection. For infection, virus stocks were diluted in OptiPRO SFM (ThermoFisher Scientific) serum-free medium according to the desired MOI. For virus adsorption 200 μl (24-well) or 1 ml (6-well) of virus master mix was added to the cells and incubated at 37°C in a 5% CO2 atmosphere with 95% humidity. After one hour, virus dilutions were removed, cells were washed three times with PBS and wells were refilled with DMEM infection medium. To determine infectious titers, supernatants were harvested at the indicated time points, and diluted 1:2 in virus preservation medium and stored at −80°C until conducting plaque titration assay.

### Infection bronchial epithelial cells

Human bronchial airway epithelial (hBAE, SmallAir^TM^) cell cultures applied in Figure 2B were obtained from Epithelix Sàrl, Geneva Switzerland. All other experiments were conducted with hBAECs isolated from explanted lungs which were obtained from the Hannover Lung Transplant Program after patients informed consent, ethical vote 2923-2015. For isolation of hBAECs, human bronchial tissue was cut into small pieces in Hank’s buffer (Thermo Fisher Scientific) containing 0.18% protease XIV and incubated for two hours at 37°C. After thorough pipetting with a 25/50 ml serological pipette, cell solution was filtered through a 100 µm cell strainer (Corning) to remove clumps and 10 ml RPMI supplemented with 10% FCS (Thermo Fisher Scientific) was added. After centrifugation for 10 min at 500*g* and 4°C, supernatant was removed and cells were resuspended in SAGM^TM^ (Lonza) + Primocin (InvivoGen) + Penicillin-Streptomycin (P/S) (Sigma-Aldrich). For air-liquid interface cultures, 200.000 hBAECs were seeded onto PureCol- (Advanced BioMatrix) coated 12 well inserts (Greiner Bio-One) in SAGM^TM^ + Primocin + P/S. 48 hours post seeding, culture medium in apical and basal chamber was changed to PneumaCult-ALI medium (STEMCELL Technologies). Air-Lift was performed 48 hours later by gently removing medium from the apical chamber. Homogenous distributed cilia were visible latest three weeks after air-lift and inserts were used for infections.

For infection, the apical surface was washed up to five times with 200 μl PBS to remove mucus. Virus stocks were diluted in OptiPRO and hBHAE were infected with an absolute infectious dose of 50,000 PFU (SmallAir^TM^) or 100,000 PFU (in-house hBAECs). Cells were incubated for 1.5 hours at 37°C in a 5% CO_2_ atmosphere with 95% humidity. After adsorption, virus dilutions were removed and the cells were washed three times with 200 μl PBS. Samples were taken at the indicated time points from the apical surface by applying 200 μl PBS to the cells. PBS was incubated on the cells for 10 minutes at 37°C to ensure that virus particles diffuse into the solution before collecting the supernatant samples. Basolateral medium (SmallAir™ Medium for SmallAir^TM^ cultures or PneumaCult-ALI for in-house hBAECs) was exchanged every 48 hours.

### Infection of nasal airway epithelial cells

Primary human nasal airway epithelial cells (hNAECs) were collected from healthy individuals by nasal brushings. Informed consent was obtained from all volunteers and the study was approved by the Charité Ethics Committee (EA2/161/20, EA2/066/20). Cultivation of hNAECs was performed as previously described (Gentzsch et al., 2017). Briefly, cells were expanded using the conditionally reprogrammed cell (CRC) culture method, then p.2 or p.3 cells were seeded on porous Transwell or Snapwell 1.1 cm^2^ supports (Corning) in UNC-ALI medium and differentiated at air-liquid interface for at least three weeks prior to infection. Approximately 200,000 hNAECs were infected with SARS-CoV-2 B.1 or B.1.1.7/v2 at an MOI of 0.1 in 150 µl D-PBS containing 0.3 % BSA for one hour at 37°C in a 5% CO_2_ atmosphere with 95% humidity. Afterwards, cells were washed apically with D-PBS and fresh medium was added basolaterally. Samples were taken at the indicated time points from the apical surface by applying 100 μl D-PBS to the cells. PBS was incubated on the cells for 30 minutes at 37°C to ensure that virus particles diffuse into the solution before collecting the supernatant samples. A 250 µl sample was taken from the basolateral side and medium was replenished. All samples were titrated on Vero E6 cells by plaque assay to determine infectious titers.

### Infection of lung organoids

Human lung organoids were established as previously published (Youk et al., 2020). Informed consent was obtained from all volunteers and the study was approved by the Charité Ethics Committee (project 451, EA2/079/13). For infection, Matrigel was liquefied and removed on ice and organoids were broken up by repeated resuspension using a disposable syringe with needle (27G). Virus stocks were diluted at the desired MOI in organoid infection medium (Advanced DMEM/F12 with 10 mM HEPES and 1x GlutaMax, ThermoFisher Scientific) and dilution was inoculated for one hour at 37°C and in 5% a humidified CO_2_ atmosphere. After infection, organoids were washed twice with PBS and resuspended in Cultrex 3-D Culture Matrix (R&D Systems) for 30 min before organoid medium (as described above) was added. Samples were taken from supernatants at the indicated time points and analyzed by plaque titration assay as described previously.

### Infection of intestinal organoids

Human normal colon organoids were established from non-cancerous parts of colorectal cancer resection tissue and cultured as previously published (Sato et al., 2011) under the ethics approval no. EA4/164/19 (to Markus Morkel). For infection studies, organoids were harvested, Matrigel (Corning, #356231) was removed by resuspension and centrifugation. Subsequently, organoids were infected in solution with an MOI of 0.05 (SARS-CoV-2 WT and B.1.1.7 strains) at 37°C for one hour. Infected organoids were seeded in Matrigel and were supplemented with medium. Samples were taken from supernatants at 24, 48 and 72 hours post-infection and analyzed by real-time RT-PCR as described (Corman et al., 2020).

### Synchronized infection experiments

Synchronized infection experiments were performed to determine entry efficiency of the virus variants. Infection of cells was performed on ice and cells were immediately transferred to 4°C for one hour after virus dilutions were added to ensure synchronized virus uptake and start of replication. After virus adsorption, cells were washed five times with PBS to remove excess of virus particles. Cells were lysed either immediately, or incubated with infection medium until four or six hours post-infection. At the indicated time points, medium was removed and cells were lysed with MagNA Pure 96 external lysis buffer (Roche, Penzberg, Germany). Isolation of RNA from cell lysates and quantitative RT-PCR on subgenomic nucleocapsid RNA was performed as described elsewhere (Corman et al., 2020; Kreye et al., 2020). Entry inhibitors were dissolved in DMSO in the indicated concentrations, added one hour prior to virus infection and were supplied for the entire duration of the experiment.

### Plaque assay

Plaque assay was performed to determine the titer of stocks and the infectious dose of supernatants harvested from infected cells. 175.000 Vero E6 cells were seeded in a 24-well plate one day prior to infection. After washing the cells once with PBS, cells were inoculated in duplicates with 200 µl of 1:10 serially diluted cell culture supernatants from infected cells. After adsorption for one hour at 37°C, the virus dilutions were removed and 500 μl of a highly viscous overlay (1:1 mix of 2.4% avicel and 2x concentrated DMEM supplemented with 5% FBS, 2% NEAA and 2% NaP) was added to each well. The overlay was discarded at three days post-infection. Cells were fixed for 30 min in 6% formaldehyde, washed once with PBS and stained for 15 min with crystal violet solution. Plaques were counted from one to two dilutions for which distinct plaques (in a range between 1-100 plaques) were detectable. To calculate the titer, the number of all plaques counted was divided by the respective inoculation volume and multiplied with the inversed dilution factor.

### Competition assay

Calu-3 cells were infected in 24-well plates with a mixture of two SARS-CoV-2 variants, using three different ratios (1:1 and 9:1 and 1:9) and an initial, total infectious dose of 10.000 PFU (corresponding to an MOI of 0.04). Serial infections were performed by sampling the supernatant of the previous passage at 24 hours post-infection and infecting naive Calu-3 cells with a 1:50 dilution of this sample. This process was repeated until completion of five passages. As a control for genome stability over five passages, single infections were performed. Viral RNA was isolated from the initial inoculum and from the supernatant of all five passages. To confirm that the virus is detectable over five passages, concentration of viral RNA was analyzed by quantitative RT-PCR (E gene assay) from each passage. To determine the variant frequency in each passage, RNA samples were sequenced using next generation sequencing techniques (Illumina technology). For virus sequence analysis, the raw sequences were trimmed, matched and presorted for SARS-CoV-2-specific sequence reads. The processed sequence reads were mapped to the BetaCoV/Munich/ChVir984/2020 genome (here referred to as SARS-CoV-2 2019-nCoV strain) in Geneious (version 9.1.8). The two virus variants in each sample were distinguished from each other by their lineage-specific mutations. For evaluation, the relative variant frequencies were calculated for each variable position. This was conducted for a total of 19 lineage-specific mutations which were distributed over the entire genome.

### Next generation sequencing

Viral RNA was extracted using MagNA Pure 96 System (Roche, Penzberg, Germany) according to the manufacturers’ recommendations. The RNA-seq library was prepared from viral RNA extracts using the KAPA RNA HyperPrep Kit (Roche, Penzberg, Germany) and KAPA DI adaptors according to the manufacturers’ instructions. The RNA library was subjected to next generation sequencing on a NextSeq System (Illumina) using a NextSeq 500/550 v2.5 Kit (Illumina). Sequences were analyzed using the geneious software, version 9.1.8, and sequence reads were assembled by mapping reads to the respective reference sequences.

### Live cell imaging

For live cell imaging, Vero E6 cells were infected with SARS-CoV-2 984, B1.1.7 Passau or B1.1.7 Baden-Württemberg at an MOI of 0.01 or 0.001 for one hour with subsequent replacement of the inoculum with full culture medium. Cells were imaged with the Zeiss LSM800 Airyscan Confocal Microscope over 72 hours with 30 min intervals in a 5% C0_2_ supplemented, humidified environment. Images were analyzed for the onset of visible cytopathic effects and merged using Zeiss ZEN Blue 3.0 and ImageJ 1.53c.

### In vivo infections

Animals: Animal procedures were performed according to the European Guidelines for Animal Studies after approval by the relevant state authority (Landesamt für Gesundheit und Soziales, Berlin, permit number 0086/20). Per group, nine male and female Roborovski dwarf hamsters (*Phodopus roborovskii*) obtained via the German pet trade were used. Animals were housed in groups of three-six hamsters in GR-900 IVC cages (Tecniplast, Buguggiate, Italy) and provided with bountiful enrichment and nesting materials (Carfil, Oud-Turnhout, Belgium). Hamsters of the same sex were randomly distributed into experimental groups and individually marked with a subcutaneously implanted IPTT-300 transponder (BMDS, Seaford (DE), USA) that allows remote identification and measurement of body temperature.

Infection and transmission experiments: To determine virus production *in vivo*, nine hamsters were inoculated with 100,000 PFU of either WT or B.1.1.7. as previously described (Trimpert et al., 2020). Briefly, anaesthetized hamsters received 100,000 PFU SARS-CoV-2 in 20 µL MEM by intranasal instillation. At 24 hours post-inoculation, contact to uninfected hamsters was enabled by placing three infected animals into a cage containing three uninfected animals of the same sex. Hamsters were monitored twice daily for clinical signs of infection. Body weight and temperature was recorded daily. Hamsters were sacrificed to determine virological parameters of infection on days 2, 3, 5 and 7 post-infection or contact, or once an individual reached a defined humane endpoint.

Virus titrations, RNA extractions and RT-qPCR: To determine virus titers from 50 mg of lung tissue, tissue homogenates were prepared using a bead mill (Analytic Jena) and 10-fold serial dilutions were prepared in MEM, which were then added to Vero E6 cells in 12-well plates. The dilutions were removed after two hours and cells were overlaid with 1.25% microcrystalline cellulose (Avicel) in MEM supplemented with 10% FBS and penicillin/streptomycin. Two days later, cells were formalin-fixed, stained with crystal violet, and plaques were counted. RNA was extracted from 25 mg of lung homogenates and oropharyngeal swabs using the innuPREP Virus RNA kit (Analytic Jena). Viral RNA copies were quantified in 10% of the obtained eluate volume with a one-step RT-qPCR reaction using a standard curve and the Luna Universal Probe One-Step RT-qPCR kit (New England Biolabs) and previously published TaqMan primers and probe (Corman et al., 2020) on a StepOnePlus RealTime PCR System (Thermo Fisher Scientific).

### Reverse genetics

We employed the previously described in-yeast transformation-associated recombination (TAR) cloning method (Thi Nhu Thao et al., 2020) for the generation of infectious SARS-CoV-2 B.1.1.7 cDNA clones. Overlapping DNA fragments were obtained by first strand cDNA synthesis of viral RNA extracts from infected Vero E6 cells using SuperScript III reverse transcriptase (Invitrogen) followed by a nested Phusion PCR (Invitrogen). Primers for TAR fragment generation were used as previously described (Thi Nhu Thao et al., 2020) with B.1.1.7-specific deviations for two fragments, as specified in Table S1. For generation of the D^614^G mutant we performed site-directed mutagenesis PCR (NEB) on synthetic viral subgenomic fragments cloned into pUC57 vectors (Thi Nhu Thao et al., 2020). Assembly of purified DNA fragments was performed by TAR cloning as previously described (Thi Nhu Thao et al., 2020). Clones were screened for correctly assembled DNA fragments by multiplex PCR using the QIAGEN Multiplex PCR kit (QIAGEN) according to the manufacturers’ instructions. Clones tested positive for all junctions were expanded, plasmid DNA was extracted, linearized and subjected to T7-based *in vitro* RNA transcription (Thermo Fisher Scientific). Capped viral RNA was electroporated into baby hamster kidney cells and supernatant was subsequently transferred to Vero E6 cells one day after electroporation for stock production. Successful virus rescue was confirmed by SARS-CoV-2-specific RT-PCR. Virus stocks were harvested three days post-infection, purified and deep sequenced as described above.

### Isolation of viral RNA and quantitative real-time RT-PCR assay

For isolation of viral RNA, 50 µl of supernatant was diluted in 300 µl of MagNA Pure 96 external lysis buffer (Roche, Penzberg, Germany). All samples were heat inactivated for ten minutes at 70°C prior to export from the BSL-3. Isolation and purification of viral RNA was performed using the MagNA Pure 96 System (Roche, Penzberg, Germany) according to the manufacturers’ recommendations. Viral RNA was quantified using real-time RT-PCR (E gene assay) as previously described (Corman et al., 2020).

### Isolation of total RNA, cDNA synthesis and quantitative PCR

For extraction of total RNA, the MagNa Pure 96 System (Roche, Penzberg, Germany) was used according to the manufacturers’ instructions. Briefly, cells were washed once with PBS before 350 µl of external lysis buffer (Roche, Penzberg, Germany) was added to the cells. Lysed cells were resuspended 2-3 times and transferred to the reaction tube. Samples were heat-inactivated for ten min at 70°C and exported from the BSL-3 laboratory. For quantitative RT-PCR, a 12.5 µl reaction with 2.5 µl RNA was done with the SuperScript III one-step reverse transcriptase-PCR system (Invitrogen) with the Platinum Taq DNA polymerase according to the manufacturers’ protocol and the primers indicated in Table S2. Probes contained a 5’ FAM-520 reporter dye and a ZEN/Iowa Black FQ 3’ quencher (Integrated DNA technologies). The RT-PCR was performed using a thermocycling protocol with reverse transcription for 15 min at 55°C and a subsequent denaturation step for two min at 95°C to restore *Taq* DNA polymerase activity, followed by PCR amplification by 45 cycles of 95°C for 15 sec and 58°C for 30 sec. Fluorescence signals were detected after the elongation step of each cycle. The mean fold change in gene expression was calculated by the delta-delta ct method and by using expression of TATA-binding protein (TBP) as a reference.

To determine early virus replication, a quantitative RT-PCR targeting the subgenomic RNA encoding the nucleocapsid (sgN) was performed. Viral RNA was extracted from cell lysates which were previously lysed by external lysis buffer (Roche, Penzberg, Germany) as described above. RT-PCR was done with the following primers and probe: nCoV sgN Fwd: 5’-CGA TCT CTT GTA GAT CTG TTC TC-3’, nCoV sgN Rev: 5’-CAG TAT TAT TGG GTA AAC CTT GG-3’ and nCoV sgN prb: 5’-56-FAM/ CAG TAA CCA GAA TGG AGA ACG CAG /3BHQ-1-3 ((Kreye et al., 2020))’. For quantification, values were normalized to the housekeeping gene TBP levels by delta ct method.

### Tat-mediated cell-cell fusion assay

CHO and TZM-bl cells were retrieved from the NIH AIDS Reagent Program and propagated as recommended. CHO cells were transiently transfected with expression plasmids for HIV-1 Tat and individual pCG-spike-HA or empty vector control for 48 hours, using Lipofectamine LTX Reagent with PLUS™ Reagent (Invitrogen). TZM-bl cells, stably expressing LTR-driven luciferase, were transfected with a plasmid encoding human ACE2 and human myc-TMPRSS2. CHO and TZM-bl cells were cocultured for eight hours. Subsequently, cells were washed once with PBS, lysed using cell culture lysis buffer (Promega), and Tat-dependent increase of luciferase enzyme activity in cell lysates was determined with the Luciferase Assay system (Promega). Luminometric activity was analyzed with a Mithras luminometer.

### Lentivirus production and transduction experiments

SARS-CoV-2 spike-HA-pseudotyped lentiviral particles were produced in triple-transfected HEK293T cells. Cells were transfected with individual pCG-SARS-CoV-2 spike-HA plasmids, the HIV-1-based packaging plasmid deltaR8.91 (Zufferey et al., 1997) and the luciferase transfer plasmid pCSII-luciferase (Agarwal et al., 2006) via calcium phosphate precipitation. Virus-containing supernatant was harvested 40 and 64 hours post transfection and sterile-filtered. Particles were concentrated via ultracentrifugation through a 20% sucrose cushion. Indicated cell lines were transduced for 72 hours with identical p24 capsid equivalents as quantified by immunoblotting of particle lysates. Transduction efficiency was quantified luminometrically three days post-transduction.

### Immunoblotting

To determine incorporation and processing of Spike in lentiviral particles, transduced cells and lentiviral particles were lysed with M-PER Mammalian Protein Extraction Reagent (Pierce) and Triton X-100, respectively. The lysate was mixed with Laemmli buffer and boiled for ten minutes at 95°C. Proteins were separated on a 10% SDS-PAGE and immobilized on a nitrocellulose membrane (GE Healthcare) using the Trans-Blot Turbo system (BioRad). Blocked membranes were incubated with the following antibodies: mouse anti-HIV-1 p24 capsid (ExBio, 1:1000), rabbit anti-S2 spike (Novusbio, NB100-56578, 1:1000), mouse anti-HA (Sigma, H3663, 1:1400), rabbit anti-tubulin (Cell Signaling Technology, 2144S, 1:1000). Secondary antibodies conjugated to Alexa680/800 fluorescent dyes were used for detection and quantification by Odyssey Infrared Imaging System (LI-COR Biosciences). Spike processing efficiency was calculated as the percentage of S2 from total spike signal. Relative levels of spike-HA abundance in lentiviral pseudotypes were quantified by calculating the signal intensity of S2-HA per HIV-1 p24 capsid.

To determine the processing and incorporation of spike from infected cells, cells and purified virus particles were lysed with RIPA (ThermoFisher Scientific) buffer supplemented with complete protease inhibitor cocktail (Roche) for 30 min at 4°C. Subsequently, lysates were centrifuged for 15 min at 4°C and 15,000 rpm to remove cell debris. The supernatants were mixed with 4x Laemmli buffer, which was supplemented with 10% beta-mercaptoethanol, and lysates were boiled for ten minutes at 95°C to ensure protein denaturation and virus inactivation. Protein concentration was determined by BCA protein assay (ThermoFisher Scientific) and 20 µg total protein was loaded. Proteins were separated by SDS-PAGE on a 6% gel and transferred to a nitrocellulose membrane (0.45 µm pore size, GE Healthcare) by Trans-Blot Turbo system (BioRad). Membranes were blocked with 5% dried milk in 0.1% PBS-Tween (0.9% NaCl, 10 mM Tris-HCl [pH 7.5], 0.1% Tween 20) for 30 min at room temperature. Blocked membranes were incubated with the following antibodies: rabbit anti-S2 spike (Novusbio, NB100-56578, 1:1000), rabbit anti-SARS-CoV-2 nucleocapsid (GeneTex, GTX135361, 1:1000). Secondary antibodies conjugated with horseradish peroxidase (HRP) were used for chemiluminescence-based detection by Fusion Fx7 (Peqlab Biotechnologie GmbH). Detection was performed using SuperSignal™ West Femto substrate (ThermoFisher Scientific). Quantification was done by the use of ImageJ 1.48v software. Spike processing efficiency was calculated as the percentage of S2 from total spike signal. Relative levels of spike abundance in concentrated virion preparations were quantified by calculating the signal intensity of S2 per nucleocapsid.

### MagPix Luminex

To assay cytokine levels in airway epithelial cell supernatant, 25 µl of supernatant were sampled prior to infection and at 24 h and 48 h post-infection with SARS-CoV-2 WT, B.1.1.7/v1, and B.1.1.7/v2. Cytokine quantification was performed using a Human Cytokine/Chemokine/Growth Factor Panel A 48-Plex Premixed Magnetic Bead Multiplex Assay (Merck Millipore), using the Luminex MAGPIX System in 96-well plate format, according to the manufacturer’s instructions. Plate washing steps were performed using HydroFlex Microplate Washer (Tecan). Calibration, verification, and quality control checks were met for all of the analytes. However, Analyte 15 (FGF-2) was omitted because its standard curve had an R^2^ value of 0.82 and Analyte 53 (IL-17F) was omitted because it had a high limit of detection and several extreme outliers from the rest of the dataset. All other analytes were reported.

### PRNT assays

Plaque reduction neutralization tests (PRNT) were performed as previously described (Kreye et al., 2020; Wölfel et al., 2020). Briefly, heat-inactivated sera were serially diluted starting at 1:40 in OptiPro, mixed 1:1 with 200 μL virus solution containing 200 plaque forming units of SARS-CoV-2 (strains B.1, B.1.1.7/v1, and B.1.351) and 200 µL of the mix were incubated in duplicates on Vero E6 cells (160,000 cells per well) seeded in 24-well plates on the previous day. After one hour incubation at 37°C, the supernatant was discarded and cells were washed with PBS and overlaid with 1.2% Avicel solution in supplemented DMEM. After three days at 37°C, the supernatants were removed and cells were inactivated and fixed with a 6% formaldehyde/PBS solution and stained with crystal violet. Serum dilutions with a mean plaque reduction of 50% and 90% are referred to as PRNT_50_ or PRNT_90_. For numerical calculations, titers <40 were set to 20, and titers >1:640 were set to 1:1,280.

### Surrogate neutralization assay

Neutralizing capacity of patients’ sera against B.1, B.1.1.7, and B.1.351 was assessed by a surrogate virus neutralization test (cPass Assay, Medac, Wedel, Germany) as described previously (Momsen Reincke et al., 2021; von Rhein et al., 2021). Briefly, sera of infected and vaccinated patients were diluted 1:10 with sample dilution buffer, mixed 1:1 with B.1-HRP-RBD, B.1.1.7-HRP-RBD, and B.1.351-HRP-RBD (provided by Medac, Wedel, Germany) solution and incubated at 37°C for 30 minutes. Afterwards, the mixture was added to the hACE2-coated plate and incubated at 37°C for 15 minutes. After washing, 3’3,5,5-tetramethylbenzidine solution was added, and the plate was incubated in the dark at room temperature for 15 minutes. Stop solution was then added and the optical density at 450 nm was measured using a Tecan Infinite 200 PRO plate reader. For calculation of the relative inhibition of ACE2/RBD binding, the following formula was applied: Inhibition score (%) = (1 − OD value sample/OD value negative control) × 100%. Values below zero were set to zero.

## Data Presentation and Statistical Analysis

If not stated otherwise, bars and symbols show the arithmetic mean of the indicated amount of independent replicates. Error bars indicate S.D. from at least three or S.E.M. from the indicated amount of individual experiments. Statistical analysis was performed with GraphPad Prism (V 8.3.0 or 9.1.2) using two-tailed unpaired Student’s t-tests or for comparing neutralizing activities the Friedman test and Dunn’s multiple comparison unless indicated differently. *P* values <0.05 were considered significant (*), <0.01 (**), <0.001 (***), <0.0001 (****); n.s. = not significant (≥0.05).

## ACKNOWLEDGEMENTS

We thank Stefan Pöhlmann for providing the plasmids encoding SARS-CoV-2 spike-HA, ACE2, myc-TMPRSS2 and Medac GmbH, Wedel, Germany for providing HRP-VOC RBDs. We thank the NIH Reagent Program for providing critical reagents. We thank the Hannover Lung Transplant Program, Prof. D. Jonigk, Department of Pathology Hannover Medical School for providing human bronchial tissue. VMC is a participant of the Charité Clinician Scientist program funded by Charité – Universitätsmedizin Berlin and the Berlin Institute of Health. JK is supported by the Center of Infection Biology and Immunity (ZIBI). Part of this work was supported by the Bundesministerium für Bildung und Forschung (BMBF) through the projects RAPID-2 (01KI2006A) to CD, and Grant 01KI2006F (RAPID-2) to TW, RECAST (01IK20337) to MAM, VMC; NaFoUniMedCovid19, Organo-Strat (01KX2021) to CG, CD, ACH, AB, MAM and TW; VARIPath (01KI2021) to VMC by the German Ministry of Health (Konsiliarlabor für Coronaviren and SeCoV) to CD and VMC, and by the Deutsche Forschungsgemeinschaft (DFG) (SFB TR84 to CD, MAM, TW, ACH; SPP 1923, GO2153/4 grant to CG; SFB 1449 Z02 to MAM; OS143/16-1 to KO); by funding from the Einstein Foundation (EC3R) to ACH; by funding from the Berlin Institute of Health (BIH) (to CG) and Freie Universität Berlin to JT and KO. SC and MW were supported by the Goethe-Corona-Fund of the Goethe University Frankfurt. MCJ, RO and UM were supported by COFONI 3FT21, BREATH (Biomedical Research In Endstage And Obstructive Lung Disease Hannover; DZL 82DZL002A1, 82DZL002B1R2N German Centre for Lung Research (DZL), R2N, Federal State of Lower Saxony (74ZN1574).

## AUTHOR CONTRIBUTIONS

Conceptualization: DN, CG, CD
Methodology: DN, KF, SSch, FWe, JT, CG, CD, AB, MAM
Investigation: DN, KF, SSch, FWe, JT, AR, SSt, JJ, JE, JK, FP, LMJ, RO, MCJ, BT, JP, JH, FWa, MLS, NH, EMB, TV, MB, AB, JS, CM, MAM
Resources: KH, MW, TTNT, SC, LGH, UM, MM, MAMü, CG, CD, AB, MAM, LH, VMC
Writing – Original Draft: DN, CG, CD
Writing – Review & Editing: DN, AB, MAM, KO, MaMü, VMC, CG, CD
Visualization: DN, SSchr, KF, JT, AR, SSt, JE, JK, FP, LMJ
Supervision: DN, SC, MAM, ACH, VT, KO, TW, UM, MAMü, VMC, CG, CD
Funding Acquisition: AB, MAM, ACH, TW, VMC, CG, CD

## SUPPLEMENTAL INFORMATION

**Figure S1.**
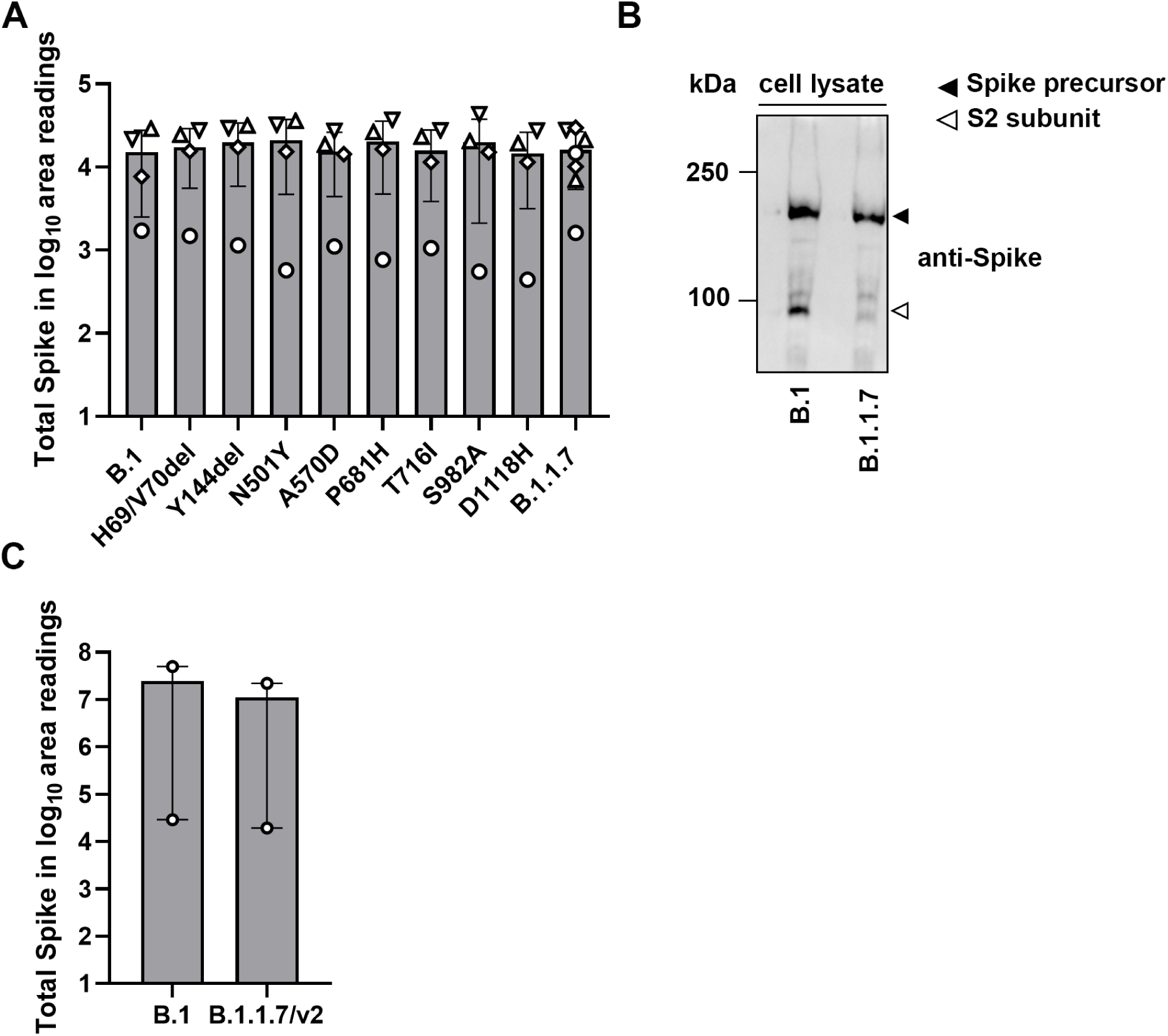
B.1 and B.1.1.7 spike is expressed at similar levels. (A) Expression of total spike in HEK-293T cells. Symbols represent independently performed experiments. (B) Vero E6 cells were infected with SARS-CoV-2 (MOI 5). Cells and virus-containing supernatants were harvested at 48 hours post-infection and processed for detection of spike by immunoblotting. (C) Expression of total Spike in Vero E6 cells was quantified by the use of ImageJ 1.48v.

**Figure S2.**
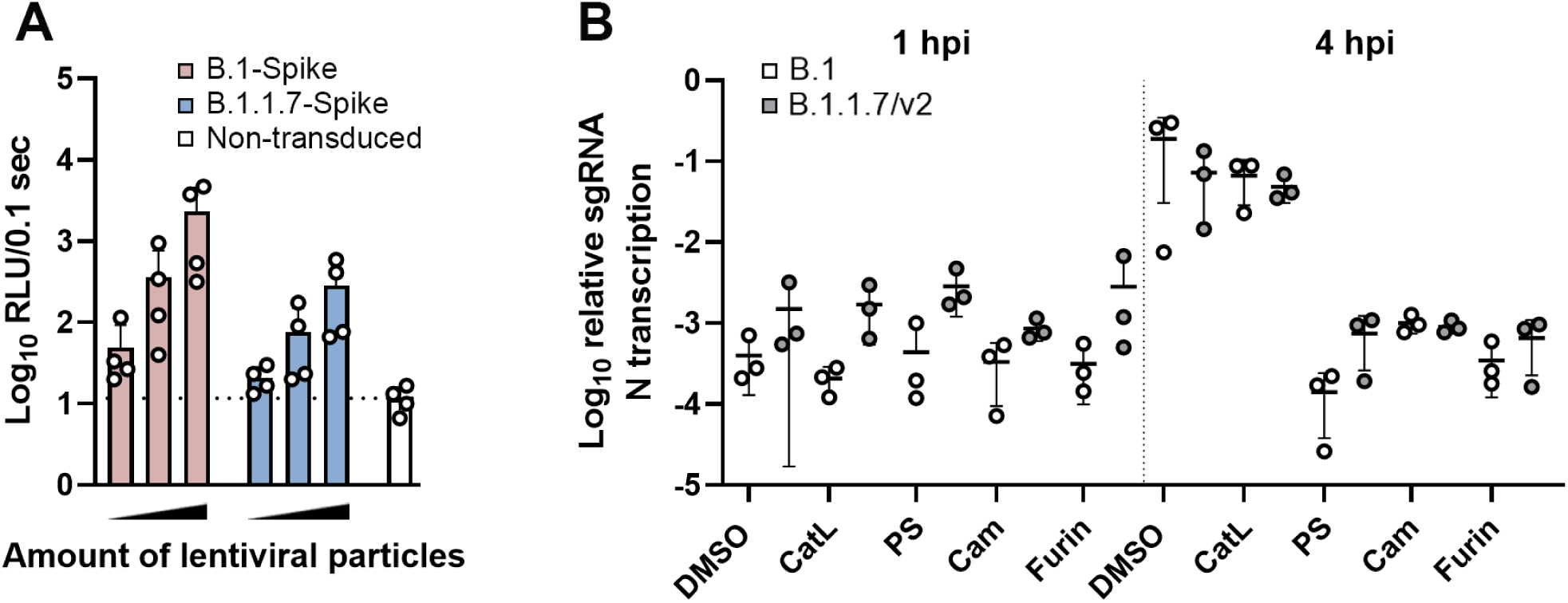
B.1.1.7 spike is not superior in mediating entry compared to B.1 spike. (A) Calu-3 cells were transduced for 72 hours with increasing amounts of lentiviral particles (0.1 µl, 1 µl and 10 µl) pseudotyped with either WT- or B.1.1.7-spike proteins. Pseudotype entry was analyzed luminometrically in cell lysates. (B) Calu-3 cells were pretreated with 25 µM MDL28170 (Cathepsin L inhibitor), 25 µM pitstop II (clathrin inhibitor), 100 µM Camostat (TMPRSS2 inhibitor) or 15 µM CMK (furin inhibitor), infected and entry efficiency was determined by sgN quantitative RT-PCR. DMSO: Dimethylsulfoxid, CatL: Cathepsin L, PS: PitStop, Cam: Camostat mesylate

**Figure S3.**
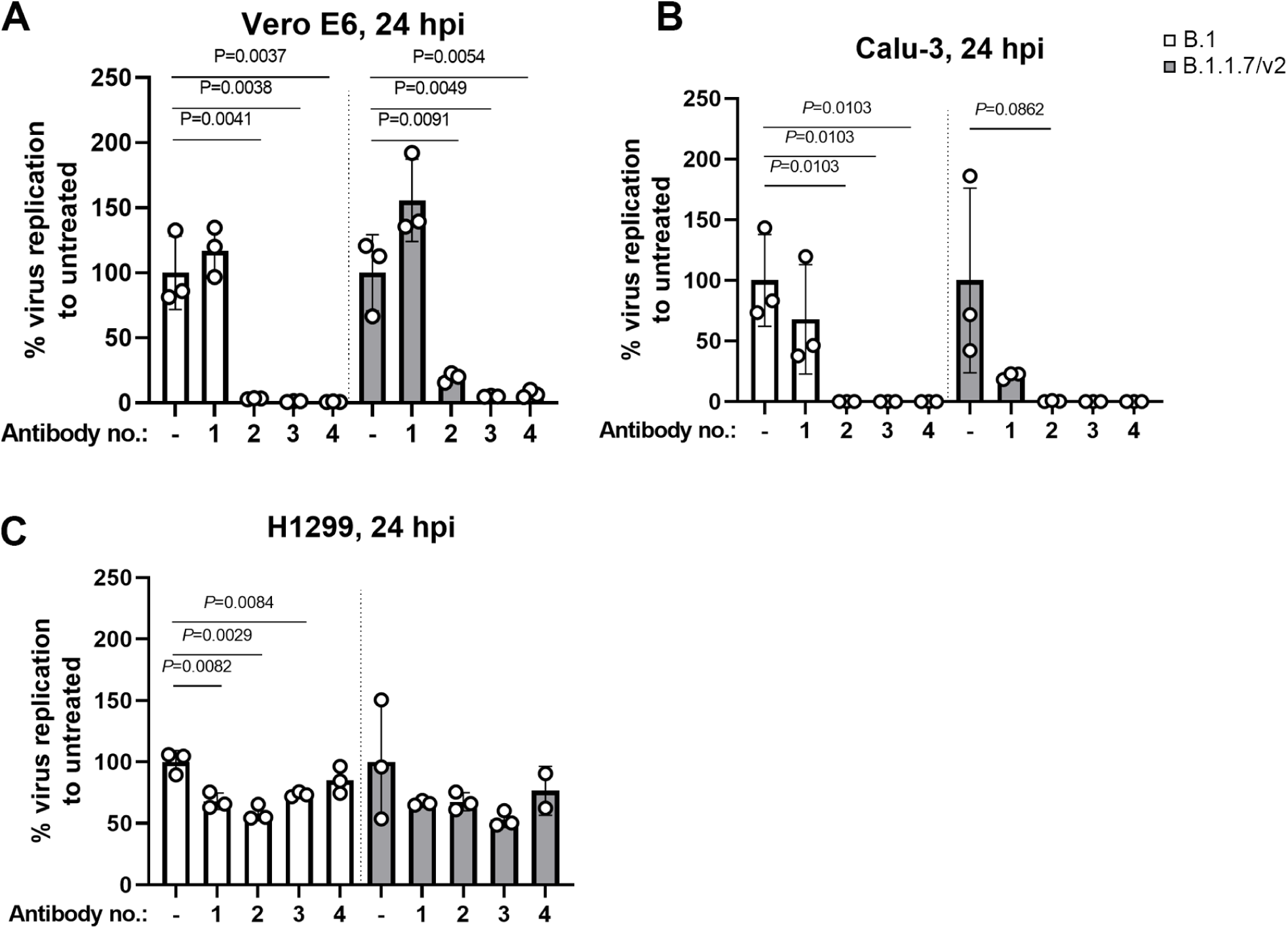
ACE2-dependent growth of SARS-CoV-2 in Vero E6 and Calu-3 cells, but not in NCI-H1299 cells. (A-C) Vero E6 (A), Calu-3 (B) and NCI-H1299 (C) cells were pretreated with four different anti-ACE2 antibodies (each applied at final concentration of 20 µg/ml) for one hour prior to infection with B.1- and B.1.1.7 isolates (MOI of 0.01). At 24 hours post-infection, viral replication was quantified from the supernatant by the use of E-gene assay. Replication was normalized to the respective untreated cells. Results from one experiment, conducted in triplicates, are shown.

## Supplemental Tables

**Table S1.**
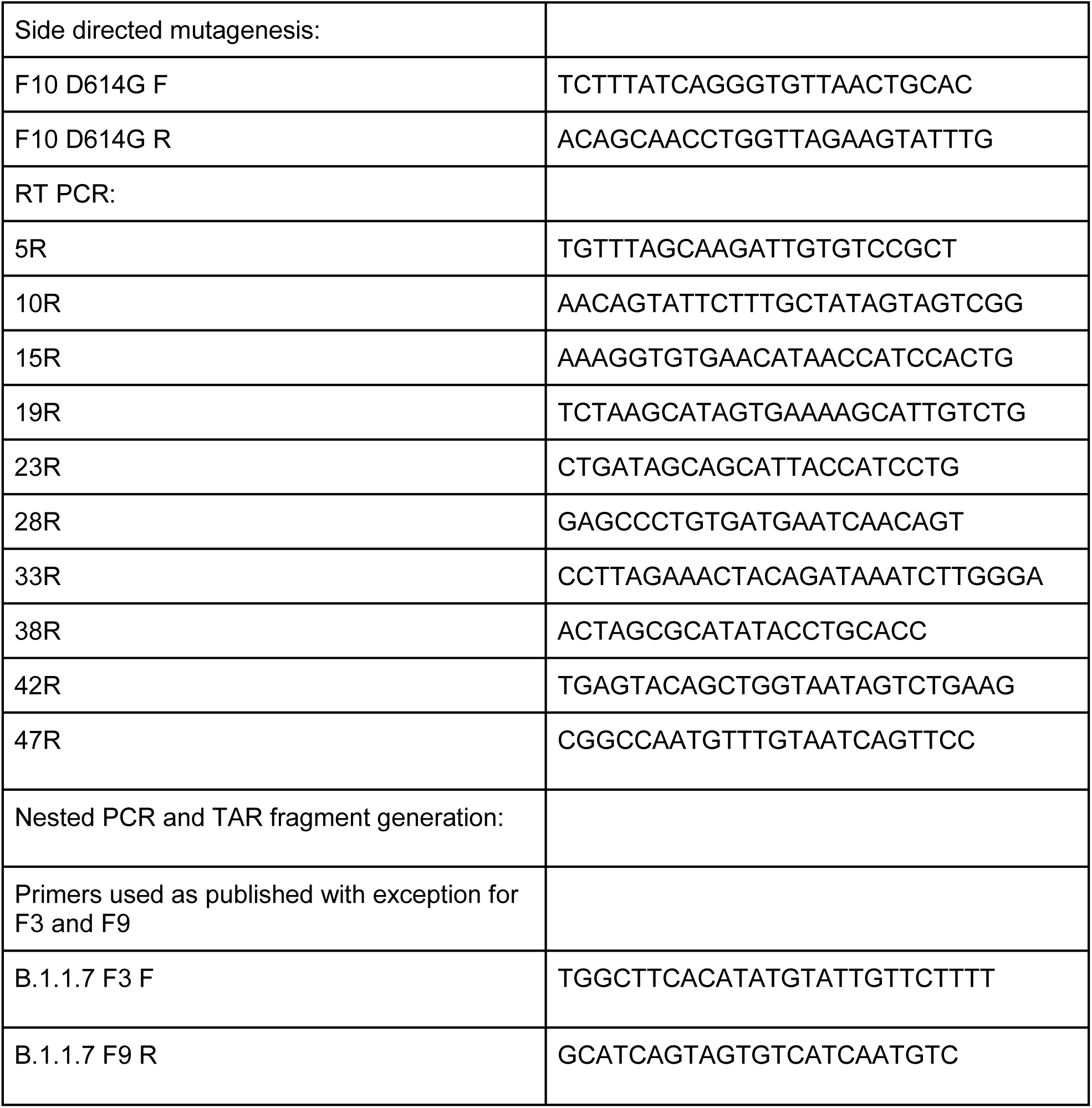
Primers used for reconstruction of recombinant SARS-CoV-2.

**Table S2.**
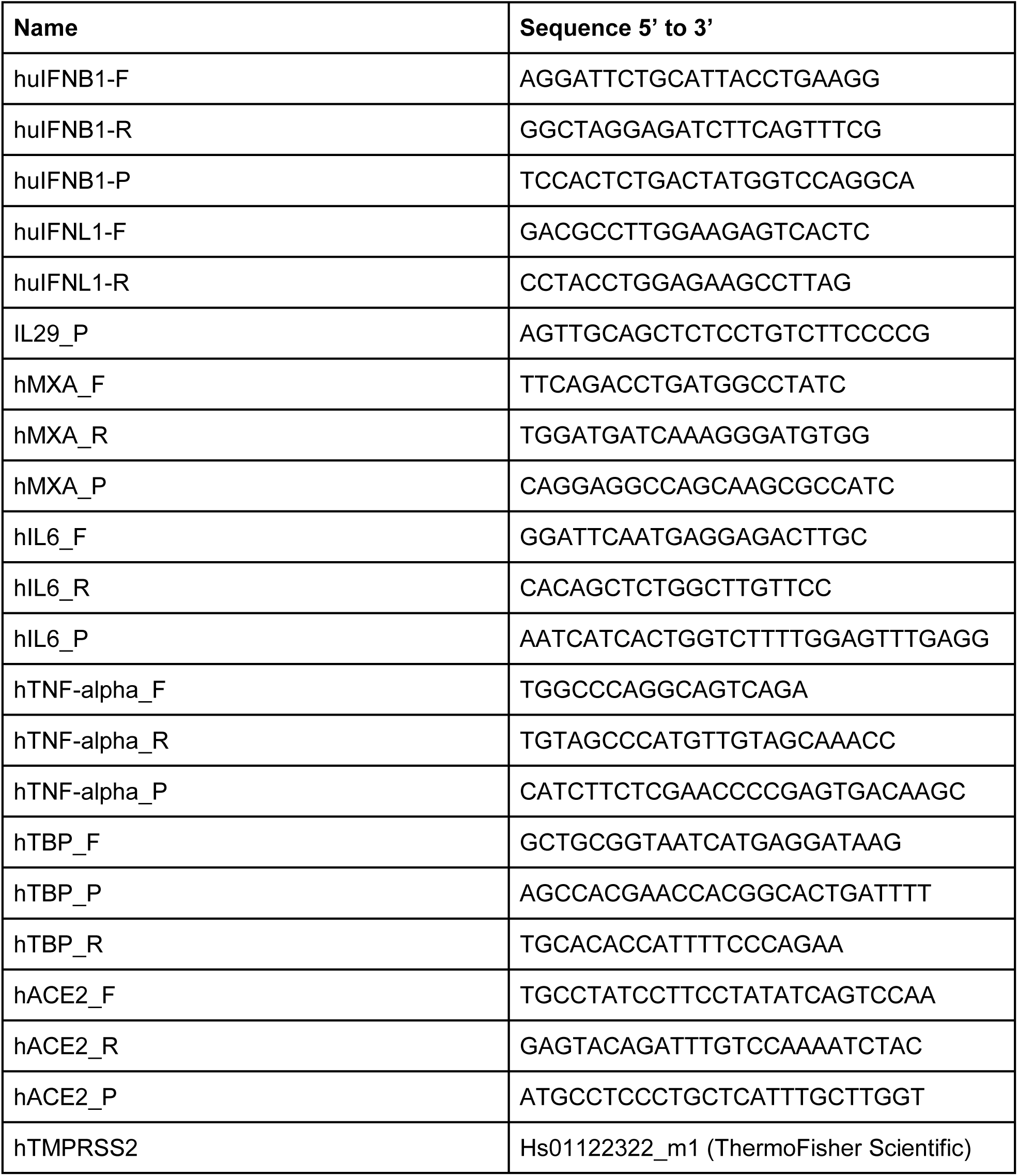
Primers used for quantitative RT-PCR.

**Supplemental Movies 1 and 2. Delayed cytopathic onset of B.1.1.7 SARS-CoV-2 infection**

Vero E6 cells were infected with B.1, B.1.1.7/v1 and B.1.1.7/v2 (MOI 0.01, supplemental movie 1; MOI 0.001, supplemental movie 2). Onset of CPE was monitored by live cell imaging until 70 hours post infection.

